# Transcript Isoform Diversity of Ampliconic Genes on the Y Chromosome of Great Apes

**DOI:** 10.1101/2023.03.02.530874

**Authors:** Marta Tomaszkiewicz, Kristoffer Sahlin, Paul Medvedev, Kateryna D. Makova

## Abstract

Y-chromosomal Ampliconic Genes (YAGs) are important for male fertility, as they encode proteins functioning in spermatogenesis. The variation in copy number and expression levels of these multicopy gene families has been recently studied in great apes, however, the diversity of splicing variants remains unexplored. Here we deciphered the sequences of polyadenylated transcripts of all nine YAG families (*BPY2*, *CDY*, *DAZ*, *HSFY*, *PRY*, *RBMY*, *TSPY*, *VCY*, and *XKRY*) from testis samples of six great ape species (human, chimpanzee, bonobo, gorilla, Bornean orangutan, and Sumatran orangutan). To achieve this, we enriched YAG transcripts with capture-probe hybridization and sequenced them with long (Pacific Biosciences) reads. Our analysis of this dataset resulted in several findings. First, we uncovered a high diversity of YAG transcripts across great apes. Second, we observed evolutionarily conserved alternative splicing patterns for most YAG families except for *BPY2* and *PRY*. Our results suggest that *BPY2* transcripts and predicted proteins in several great ape species (bonobo and the two orangutans) have independent evolutionary origins and are not homologous to human reference transcripts and proteins. In contrast, our results suggest that the *PRY* gene family, having the highest representation of transcripts without open reading frames, has been undergoing pseudogenization. Third, even though we have identified many species-specific protein-coding YAG transcripts, we have not detected any signatures of positive selection. Overall, our work illuminates the YAG isoform landscape and its evolutionary history, and provides a genomic resource for future functional studies focusing on infertility phenotypes in humans and critically endangered great apes.

## INTRODUCTION

Ampliconic Genes on the Y chromosome (YAGs) are expressed exclusively in testis (Skaletsky et al. 2003), encode proteins functioning in spermatogenesis, and are important for male fertility. In humans, YAGs belong to nine multicopy gene families—*BPY2*, *CDY*, *DAZ*, *HSFY*, *PRY*, *RBMY*, *TSPY*, *VCY*, and *XKRY*. Most of these gene families are primate-specific (Cortez et al. 2014), however some were pseudogenized or completely deleted in certain great ape lineages, which is potentially linked to interspecies differences in mating patterns (Hughes et al. 2010; Cechova et al. 2020). Although the variability in copy number (Tomaszkiewicz et al. 2016; Oetjens et al. 2016; Ye et al. 2018; Vegesna et al. 2020) and in gene expression levels (Fagerberg et al. 2014; Vegesna et al. 2019, 2020) of YAGs has been recently explored in great apes, we still do not have a full understanding of whether this variability is reflected in transcript diversity. In fact, sequences of full-length transcripts for YAG families across great apes are lacking. Moreover, their precise cellular functions in the male germline are still not completely deciphered (Lahn and Page 1997; Zou et al. 2003; Wong et al. 2004; Stouffs 2004; Yen 2004; Navarro-Costa 2012). Investigating the transcriptional landscape of YAGs is expected to inform the diversity in sperm characteristics and male fertility phenotypes observed across great apes.

Recently, great ape YAG transcripts were assembled from short-read RNA-seq data (Vegesna et al. 2020). However, only one consensus transcript was generated for most gene families and, in some cases, even that representative sequence was incomplete. These limitations arise from the fact that many YAG families are expressed at low levels (Fagerberg et al. 2014; Vegesna et al. 2019, 2020) leading to a small number of sequencing reads and problematic transcript assemblies. In another study (Cortez et al. 2014), gorilla YAG transcript sequences were reconstructed from short-read RNA-seq data originating from testis of one individual, which resulted in only one transcript per gene family. In the same study, orangutan transcript sequences for YAG families were predicted from whole-genome DNA sequencing data. The annotation of the published chimpanzee Y chromosome is incomplete, missing full-length coding sequences of all six YAG families present in this species (Hughes et al. 2010). Therefore, the complete repertoire of transcript sequences of great ape YAGs resolved at a nucleotide level is still unavailable.

To fill this critical gap, we captured (with hybridization) polyadenylated YAG transcripts from testis samples of six great ape species: human, chimpanzee, bonobo, gorilla, Sumatran orangutan, and Bornean orangutan, and sequenced them using Pacific Biosciences (PacBio) single-molecule long-read sequencing (Iso-Seq) protocol. Obtaining this unique dataset allowed us to compare full-length YAG transcripts across great ape species that diverged from each other between ~0.4 and ~13 million years ago (Locke et al. 2011; Glazko 2003). Specifically, we asked the following questions: (i) How diverse are YAG transcript isoforms across great apes? (ii) Do non-human YAG transcripts share splicing patterns with the well-annotated YAGs on the human Y chromosome? (iii) Have any transcripts accumulated nonsense mutations, leading to pseudogenization? (iv) Do any of the YAGs conserved across great ape species evolve under purifying selection? (v) Which YAG isoforms are lineage-specific, and which exhibit signatures of positive selection? By addressing these questions, we deciphered the isoform landscape and evolutionary history of YAG sequences in great apes. The resulting data and the analyses described herein will be critical for designing evidence-based strategies for preserving reproductive success of these species, all of which (except for humans) are endangered.

## MATERIALS AND METHODS

### Samples, RNA extraction, and long-read sequencing

Two human testis samples from patients without cancer (IDs: A0119c, A014a) were provided by the Cooperative Human Tissue Network (CHTN) under Penn State IRB STUDY00005084. Two chimpanzee (*Pan troglodytes*) testis samples from individuals deceased from heart failure (IDs: 8720, 9423) were provided by the University of Texas MD Anderson Cancer Center’s Michale E. Keeling Center for Comparative Medicine and Research. One bonobo (*Pan paniscus*) (ID OR5013) and one Bornean orangutan (*Pongo pygmaeus*) (ID OR3405) testis samples were provided by San Diego Zoo Institute for Conservation Research. One western lowland gorilla (*Gorilla gorilla gorilla*) (ID 2006-0091) and one Sumatran orangutan (*Pongo abelii*) (ID 1991-0051) testis samples were provided by the Smithsonian Institution.

Total RNA was extracted from all eight testis samples (~30 mg of tissue) using the RNeasy Mini Kit following the manufacturer recommended protocol (Qiagen, USA). All samples had RIN value ≥6. Two technical replicates of cDNA were generated from each of the eight RNA samples using the SMARTer PCR cDNA Synthesis Kit (Clonetech, USA). The universal 5’ cDNA primer and a 3’ barcoded technical replicate-specific oligo(dT) primer (Integrated DNA Technologies, USA) primer were used to prime the reactions (Table S1). These cDNAs were used for a selective pulldown of YAG double-stranded cDNAs using hybridization to in-house-designed biotinylated capture probes, synthesized by IDT. More detailed descriptions of the of the hybridization capture experiments are summarized in Fig. S1, Table S2. To maximize enrichment, we performed the hybridization twice.

Enrichment and hybridization followed the PacBio protocol (https://www.pacb.com/wp-content/uploads/Procedure-Checklist-cDNA-Capture-Using-SeqCap-EZ-Libraries.pdf). Probes and primers (Table S3) used to validate male-specificity were designed in-house and synthesized by IDT. To design oligo hybridization probes and primers, we generated consensus sequences using BWA-MEM (version 0.7.10) alignments (Li 2013). Illumina RNA-Seq, flow-sorted Y, and whole-genome sequencing data from chimpanzee, bonobo, gorilla, Bornean orangutan, and Sumatran orangutan (NCBI Sequence Read Archive under accession numbers: SRR10392513–SRR10392518 and SRX7685072-SRX7685081) were aligned to the reference protein-coding sequences and visualized in Integrative Genomics Viewer (version 2.3.72) (Thorvaldsdóttir, Robinson, and Mesirov 2013). Sixteen ampliconic cDNA samples (two per each of eight samples) pooled in equimolar quantities were subsequently used for preparation of Iso-Seq libraries (one “standard”: centered around ~2kb and one “longer” enriched for longer cDNA >3kb) using the PacBio protocol (https://www.pacb.com/wp-content/uploads/Procedure-Checklist-Iso-Seq-Express-Template-Preparation-for-Sequel-and-Sequel-II-Systems.pdf). The pooled samples were sequenced using the PacBio Sequel I instrument with four SMRT cells for the “standard” library and one SMRT cell for the “longer” library.

### Analysis of long reads and transcript clustering

Long reads were analyzed according to the following workflow (Fig. S2). First, BAM files of circular consensus sequences (CCSs) were produced from the raw subreads using CCS (SMRTlink, version 6; --Polish --minPasses 3). Next, the BAM files from the five SMRT cells were merged and demultiplexed using lima (version 2.2.0, https://github.com/PacificBiosciences/barcoding; --isoseq --peek-guess), with 82% of CCSs passing quality filters. For each of the 16 resulting CCS files (two technical replicates for each of the eight samples), CCSs were classified as full-length non-chimeric reads (FLNC), based on the presence of poly(A) tails, which were subsequently removed using refine (Iso-Seq3, version 3.4.0, https://github.com/PacificBiosciences/IsoSeq; isoseq3 refine --require-polya). The resulting FLNC reads were clustered at the gene family level (separation by sequence similarity) with isONclust v0.0.6.1 (--t 1 --k 11 --w 15) (Sahlin and Medvedev 2020). Each read cluster produced by isONclust represented a gene family and was subsequently error-corrected using IsoCon (--ignore_ends_len 15) (Sahlin et al. 2018). Finally, the error-corrected clusters were annotated to gene families by aligning probe sequences (Table S2) to the transcripts (Šošić and Šikić 2017) with annotate_clusters.py (script available at https://github.com/makovalab-psu/YAG_analysis). Specifically, each probe was forward- and reverse-complement and then aligned to all of the predicted transcripts in a cluster. The probe with the lowest edit distance *E* to a transcript in a cluster was voted with weight 1/*max*(1, *E*) suggesting that the cluster belongs to the gene family for which the probe is designed. The cluster was then annotated to the gene family with the largest sum of transcript weights. We annotated a cluster rather than individual transcripts because some FLNC reads were partially missing the probe or had high edit distance to all of them. However, these reads matched other regions of longer transcripts in the same cluster.

### Identification of isoforms shared between technical replicates

The sequencing and analysis strategy described above produced two clusters of transcripts, corresponding to two cDNA technical replicates per sample for each combination of sample and gene family. We let *X* and *Y* denote the two sets of transcripts corresponding to the transcripts from each of the two replicates for a given gene family. In order to eliminate random errors introduced by the PCR amplification of cDNA and subsequent sequencing (Fig. S1), we identified transcripts that were supported by both replicates by running “isoform_similarity.py” (script available at https://github.com/makovalab-psu/YAG_analysis). The script consists of two steps.

First, we identified *supported transcripts* between *X* and *Y*, denoted by *Z*. We called a transcript *t*_1_ in one of the replicates to be supported if (a) it was an exact substring of a transcript *t*_2_ in the other replicate, and (b) *t*_2_ had a maximum 5’ difference of 100 bp and a maximum 3’ difference of 30 bp to *t*_1_. Note that a transcript could be supported by several transcripts under this definition, but it only needed at least one supporting transcript to be classified as supported. Intuitively, the supported transcripts *Z* were those that were consistently predicted between replicates.

The supported transcripts could still be redundant due to experimental variability at the ends, such as 3’-end bias caused by 3’-poly(A) tail initiation of the cDNA synthesis. Therefore, as a second step, we removed the redundancy in *Z* by merging any redundant transcripts into the longest representation using the same criteria as for determining support. Specifically, we merged transcript *t*_1_ into another transcript *t*_2_ if (1) *t*_1_was an exact substring of *t*_2_ and (2) *t*_1_ fulfilled the criteria of 100 and 30 bp maximal 5’ and 3’ offsets to *t*_2_. We processed the transcripts greedily from shortest to longest transcript and, for each transcript *t*_1_, identified if there was a longer transcript it could be merged into, according to (1) and (2). If so, then we removed *t*_1_ from the set. Whatever remained in the set after this processing was the set of merged transcripts.

After identifying supported transcripts and removing redundancy, we constructed a final set of non-redundant replicate-supported transcripts (Dataset 1), which we, for brevity, from now on refer to as *replicate-supported transcripts*. The end-offsets of 100 and 30 bp were chosen to be consistent with the parameters used by the Iso-Seq3 pipeline (https://github.com/PacificBiosciences/IsoSeq) for merging transcripts, except that we did not follow their recommendation on merging transcripts differing by any internal gaps of less than 10 bp. This is because we expect transcripts within the same gene family to be highly similar and differ by only small internal gaps, and thus we kept such transcripts as separate ones.

### Prediction of coding potential

We predicted Open Reading Frames (ORFs) for all replicate-supported transcripts using getorf (Rice, Longden, and Bleasby 2000) (Dataset 2) and retained only complete ORFs of at least 50 amino acids in length (Dataset 3).

### YAG homolog sequence similarity search across great apes

We inferred statistically significant homologs according to (Pearson 2013). Protein sequence identity could be <20%, but E-values had to be <10^-6^, and bit scores had to be >50. First, we aligned replicate-supported transcripts against human and chimpanzee Y-chromosome transcripts using BLASTN (Camacho et al. 2009) (Datasets 4 and 5). Transcripts for gorilla and Sumatran orangutan were aligned to predicted or experimentally deciphered publicly available transcript sequences for these species (Camacho et al. 2009) using BLASTN (Datasets 6 and 7, respectively). Subsequently, we narrowed down the analysis to Y-chromosome protein-coding sequences (Dataset 8). To ensure that more divergent sequences were not missed, we also aligned the predicted ORFs against human Y-chromosome proteins using BLASTP (Camacho et al. 2009) (Dataset 9).

In cases where species had missing transcripts shared between technical replicates that are homologous to human Y chromosome reference sequence, we rechecked to see whether these sequences were present in the two previous steps of the analysis: (1) before merging into replicate-supported transcripts, or (2) before clustering transcripts (raw FLNC per species). Separately, we used transcripts clustered per each technical replicate and raw FLNC sequences from each species to run BLASTN analysis (Camacho et al. 2009) against the human Y-chromosome coding sequences (Datasets 10 and 11, respectively). Additionally, we predicted ORFs from transcripts clustered per each technical replicate and raw FLNC and ran BLASTP analysis (Camacho et al. 2009) against the human Y-chromosome proteins (Datasets 12 and 13, respectively).

We also counted the number of identical protein-coding sequences between samples (Table S13, Dataset 14). To do this we converted each fasta sequence into a signature and identified identical signatures that were shared using script “fasta_to_hash_4.py” (available on GitHub).

### Alternative splicing patterns and classification of splice variants

To study alternative splicing patterns of YAGs across great apes, all replicate-supported transcripts were aligned to one human genomic copy per gene family (Dataset 15) using uLTRA (version v0.0.4) (Sahlin and Mäkinen 2021) with default parameters. This allowed us to infer the evolutionary origins of replicate-supported transcripts by comparing their exon-intron structure to that of human Y-specific reference genes.

### Selection tests

For selection tests, we aligned the homologous sequences per gene family from all great apes to the reference human and macaque (outgroup) Y-chromosome coding sequences. We used the *codeml* module of PAML (version 4.8, (Yang 2007)) to estimate nonsynonymous-to-synonymous substitution rate ratio (d_N_/d_S_) for orthologous YAGs in human, chimpanzee, bonobo, gorilla, Sumatran orangutan, Bornean orangutan, and macaque (outgroup). Protein-coding sequences of all YAG replicate-supported transcripts were aligned using CLUSTALW (Larkin et al. 2007). The phylogenies were generated with the Neighbor-Joining method (with 1,000 bootstrap replicas) as implemented in MEGAX (Kumar et al. 2018). First, for each YAG family, the one-ratio model (assuming one average d_N_/d_S_ ratio for the entire tree) was compared to the two-ratio model (assuming the branch-specific d_N_/d_S_ ratio ω_s_ is different from the background d_N_/d_S_ ratio ω_o_). Second, to test for positive selection, the model that assumed the foreground ratio ω to be fixed at 1 (neutral evolution) was compared against an alternative model with branch-specific ω > 1 (positive selection). Third, to test for negative (purifying) selection, the model assuming the foreground ratio ω to be fixed at 1 (neutral evolution) was compared against an alternative model with branch-specific ω_s_ < 1 (negative selection).

## RESULTS

### Sequencing of polyadenylated Y-chromosomal ampliconic gene transcripts in six great ape species

To study the evolution of YAG transcripts across great apes, we isolated total RNA and synthesized cDNA from testis samples for six great ape species: human (two individuals), chimpanzee (two individuals), bonobo (one individual), gorilla (one individual), Bornean orangutan (one individual), and Sumatran orangutan (one individual). For each sample, we generated two technical replicates, which we started from separate aliquots of the same RNA stock and then each aliquot was processed separately (Fig. S1). After pulling down YAG cDNAs using gene-family-specific hybridization capture probes (Table S2), two PacBio Iso-Seq libraries (one with cDNA centered around ~2 kb and another one enriched for cDNA >3 kb) were pooled together and sequenced. We identified FLNCs (see Methods; Table S4) among CCSs generated from raw subreads. There were 15,529-87,639 of CCSs and 14,586-60,648 of FLNCs per technical replicate (Table S4). We next clustered FLNCs with isONclust (Sahlin and Medvedev 2020) and assigned the clusters to their respective gene families by aligning probe sequences to the FLNCs in each cluster. Finally, we performed error correction of the clustered and gene-family-assigned FLNCs with IsoCon (Sahlin et al. 2018). This resulted in the total number of transcripts ranging from 526 to 2,302 per technical replicate (Table S5). This variation in part reflected differences in the sequencing yield per technical replicate (Table S4). To decrease differences in sequencing yield among samples, and also because biological replicates were unavailable for all the species, we removed human sample 2 and chimpanzee sample 1—the samples with the highest and the lowest average number of transcripts per technical replicate, respectively (Table S5)—from subsequent analyses. We utilized these two samples to validate some of our findings from the other samples (see below). Briefly, our subsequent analyses consisted of (1) identifying replicate-supported transcripts; (2) mapping them to one human genomic copy per each YAG family, and to the databases of human and non-human full-length transcripts and protein-coding transcripts, (3) mapping predicted proteins to human Y proteins, (4) identifying identical (among samples or species) and species-specific transcripts, and (5) testing for selection acting on protein-coding transcripts (Fig. S3).

### Replicate-supported transcripts

We identified 1,510 replicate-supported transcripts, i.e. isoforms shared between the two technical replicates (Table S7), in six samples. Each replicate-supported transcript was supported by at least two reads (in each technical replicate) with an average of 92 reads (min=2, max=3,245, med=44; all but 11 replicate-supported transcripts were supported by more than two reads). The total number of replicate-supported transcripts per sample ranged from 190 (for chimpanzee) to 315 (for Bornean orangutan), with an average of 256 (Table S6). The number of replicate-supported transcripts per gene family differed substantially among gene families and among species (Fig. 1A, Table S6). The longest transcripts belonged to the *CDY* and *DAZ* gene families with mean lengths of 1,868 bp and 1,687 bp, respectively (Fig. 1B). The longest transcript was a human *DAZ* transcript (4,041 bp) followed by a chimpanzee *DAZ* transcript (3,584 bp; Fig. 1B, Tables S8-S9). The shortest transcripts belonged to the *PRY* gene family with a mean length of 374 bp (Fig. 1B, Table S8).

**Figure 1.**
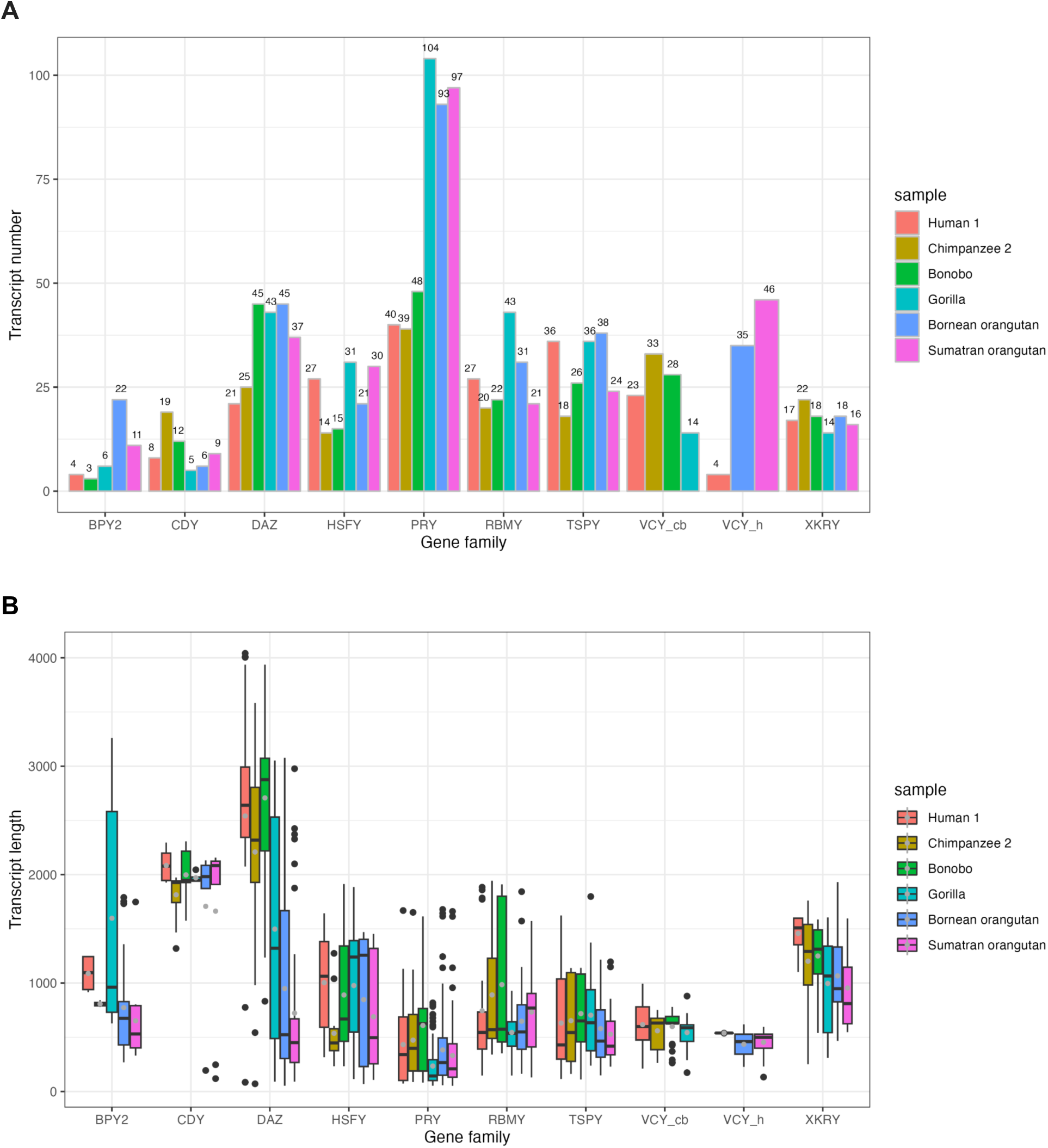
**A.** Number of replicate-supported transcripts presented separately for each great ape species and gene family. **B**. Distribution of lengths of replicate-supported transcripts per gene family per species. Black dots represent outliers, gray dots represent mean lengths, and black lines represent median lengths. *VCY_h*, captured with the human-specific probe, *VCY_cb*, captured with the chimpanzee- and bonobo-specific probe).

### Analysis of transcripts with complete open reading frames

We predicted complete open reading frames (cORFs), i.e. sequences that begin with a start codon and end with a stop codon, in the replicate-supported transcripts. Because most cORFs we found were <50 aa long (Dataset 2) we suspect them to be artifacts. However, an abundance of short proteins was found in mammalian genomes (Frith et al. 2006), therefore we have chosen an ORF length threshold of >50 aa for the downstream analysis of transcripts with cORFs (Dataset 3 and 16; Fig. 2A). In total, we identified 7,966 cORFs longer than 50 aa in our replicate-supported transcripts.

**Figure 2.**
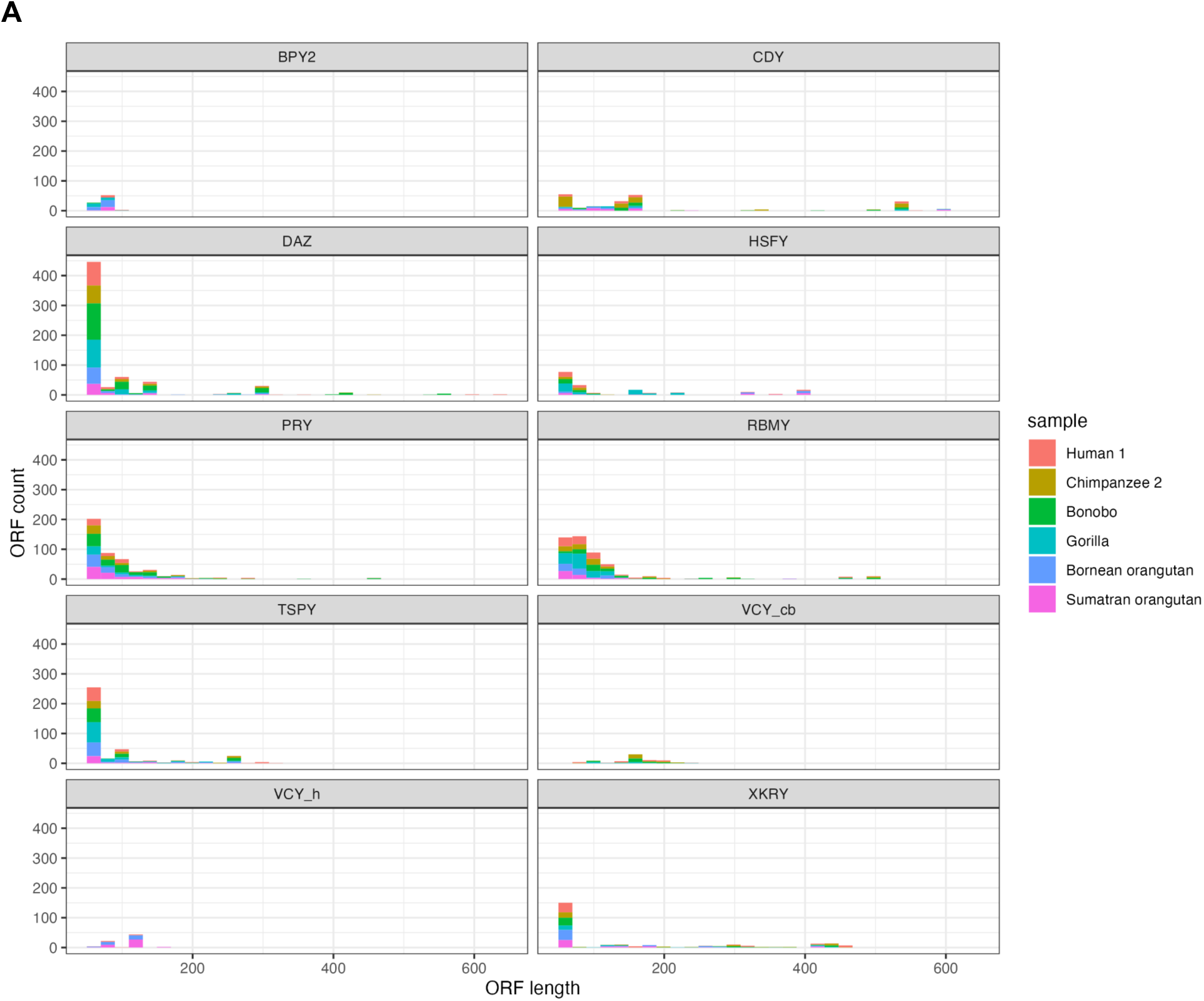
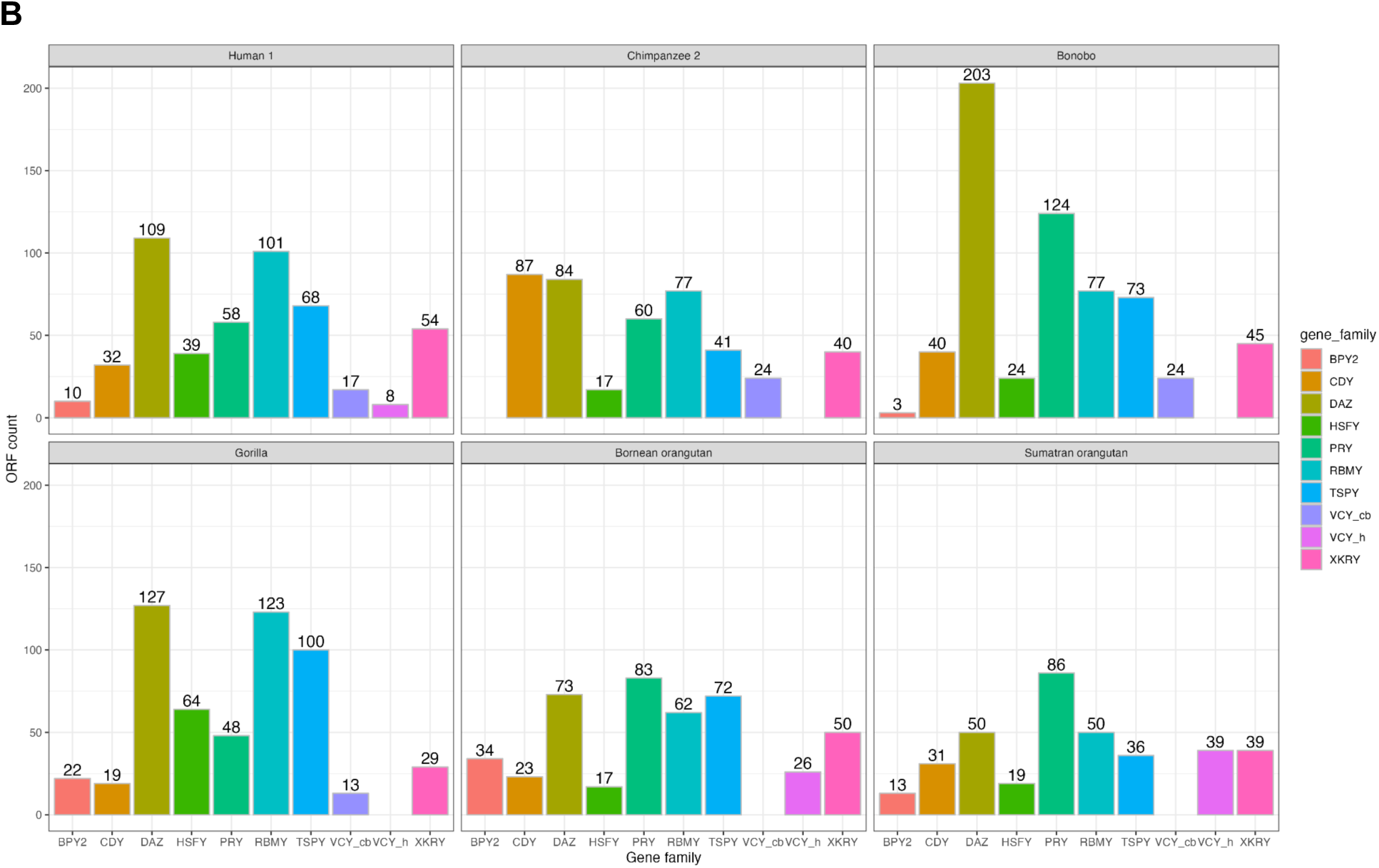
**A.** Distribution of predicted complete Open Reading Frame (cORF) lengths (in amino acids) per gene family *VCY_h*, captured with the human-specific probe, *VCY_cb*, captured with the chimpanzee- and bonobo-specific probe)—per species. **B.** Number of cORFs per gene family per species.

We recovered transcripts with cORFs from all YAG families (Fig. 2B, Dataset 16) in all the species analyzed with one exception. In chimpanzee, cORFs for all YAG families were present except for *BPY2,* whose cORF was present only in one of the technical replicates. In general, the lengths of proteins predicted from these cORFs were similar to the lengths of the corresponding human proteins reported in the literature. For example, a previous study (Lahn and Page 1997) showed human BPY2 proteins to be 106 aa long. We found cORFs corresponding to BPY2 proteins of this length in our human sample and of a similar length in our gorilla sample (Fig. 2A). The same study showed human VCY protein to be 125 aa long (Lahn and Page 2000), and we found predicted homologs of the same length in our human sample (Fig. 2A).

### Homology to human YAG protein-coding transcripts

First, to study homology to human YAG protein-coding transcripts, we used BLASTN (Camacho et al. 2009) to align all replicate-supported transcripts against publicly available Y transcript data sets for human, available in ENSEMBL (Dataset 4), chimpanzee, available in ENSEMBL (Dataset 5), gorilla Y, published by Cortez and colleagues (Cortez et al. 2014) (Dataset 6), and the predicted orangutan Y chromosome transcripts generated by Cortez and colleagues (Cortez et al. 2014; Dataset 7). Because we could not find any replicate-supported transcripts that fully covered known transcripts from the above datasets (Datasets 4, 5, 6, 7) at 100% sequence identity, we limited our subsequent analyses to protein-coding sequences.

Second, we aligned our non-human replicate-supported transcripts to the annotated human reference YAG protein-coding sequences (Dataset 8) using BLASTN (Camacho et al. 2009) (see Methods for details; Table S10). For several great ape species, we found replicate-supported transcripts homologous to human reference transcripts to be either completely absent or of reduced length in some YAG families (Table S10), suggesting deletion or pseudogenization. No homologous *BPY2* transcripts were found in chimpanzee and bonobo, however such sequences were present in one chimpanzee technical replicate. *BPY2* homologs differed substantially in length between human and gorilla (covering only 60-71% of the human reference with 98-99% of sequence identity), and even more so between each orangutan species and human (covering 34-37% of the human reference with 93-96% of sequence identity). *HSFY*, *PRY,* and *XKRY* homologs were short or absent in chimpanzee and bonobo, confirming previous findings (Bellott et al. 2014; Cechova et al. 2020; Cortez et al. 2014). *VCY* homologs were found in bonobo and in both species of orangutan, which contradicts previous results (Cortez et al. 2014; Cechova et al. 2020) and requires further investigation. *XKRY* homologs were present in gorilla (covering 80% of the human reference with 86.8% of sequence identity) and both species of orangutan (64-100% coverage with 84-93.3% of sequence identity), but were absent from both bonobo and chimpanzee (Table S10), which confirms previous studies (Hughes et al. 2010; Cechova et al. 2020).

We also aligned the predicted cORFs in replicate-confirmed transcripts to human reference Y-chromosome proteins (Dataset 9) using BLASTP (see Methods for details). The resulting high-confidence cORF homologs per gene family per species are presented in Figure 3 and Dataset 9. CDY, DAZ, RBMY, TSPY, and VCY ORF homologs were abundant across all great ape species (Fig. 3A). Among YAG families, CDY ORF homologs aligned to human proteins with the highest coverage (Fig. 3B). The largest numbers of CDY and DAZ ORF homologs were observed in bonobo. Only single HSFY ORF homolog with low coverage (17-25%) but relatively high identity (77-82%) was found in each of the chimpanzee and bonobo samples (Fig. 3A, Dataset 9). BPY2 ORF homologs were present only in human and gorilla samples. PRY homologs were missing in all species but gorilla (Fig. 3A), indicating that these sequences are either not expressed or expressed below our detection level.

**Figure 3.**
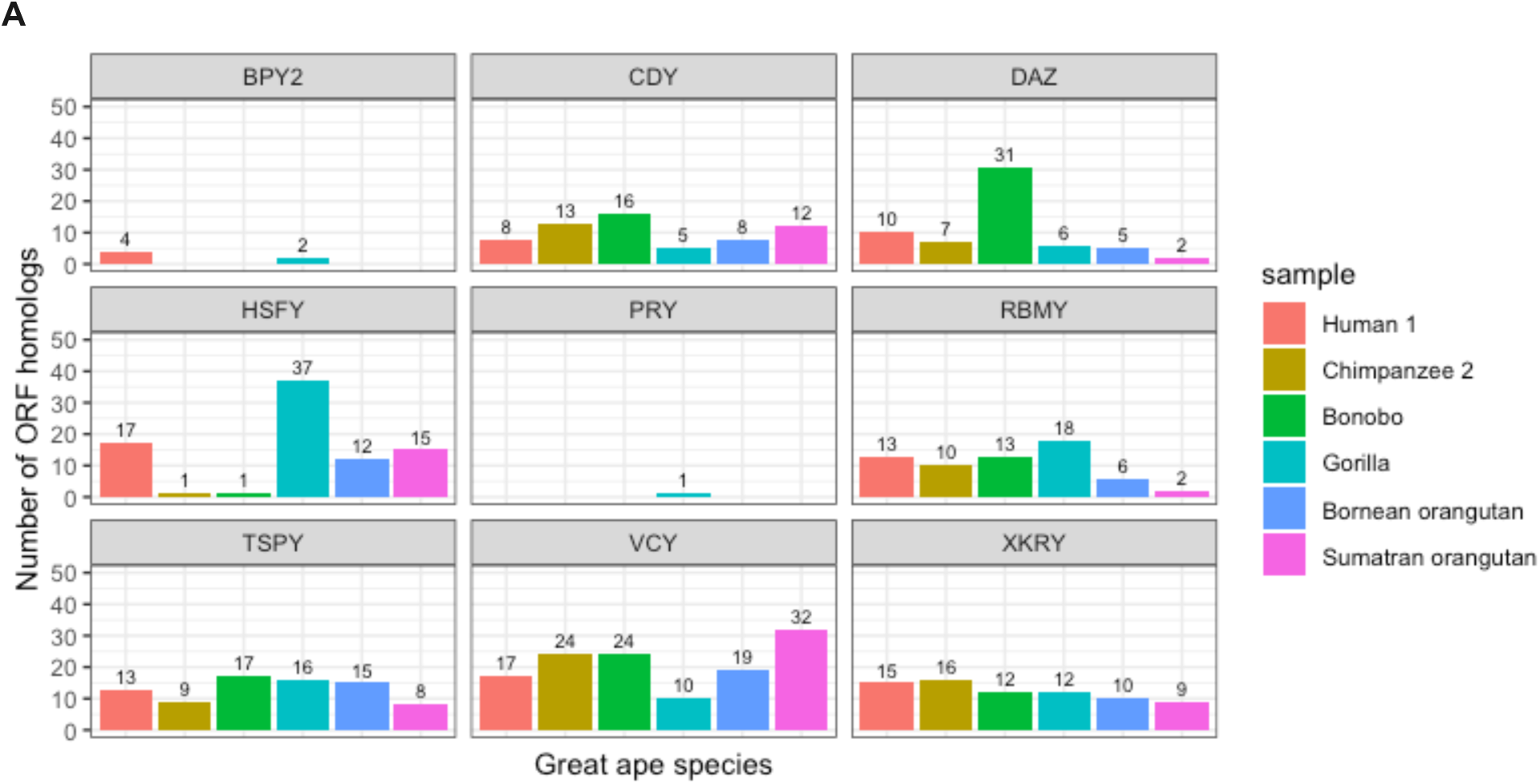
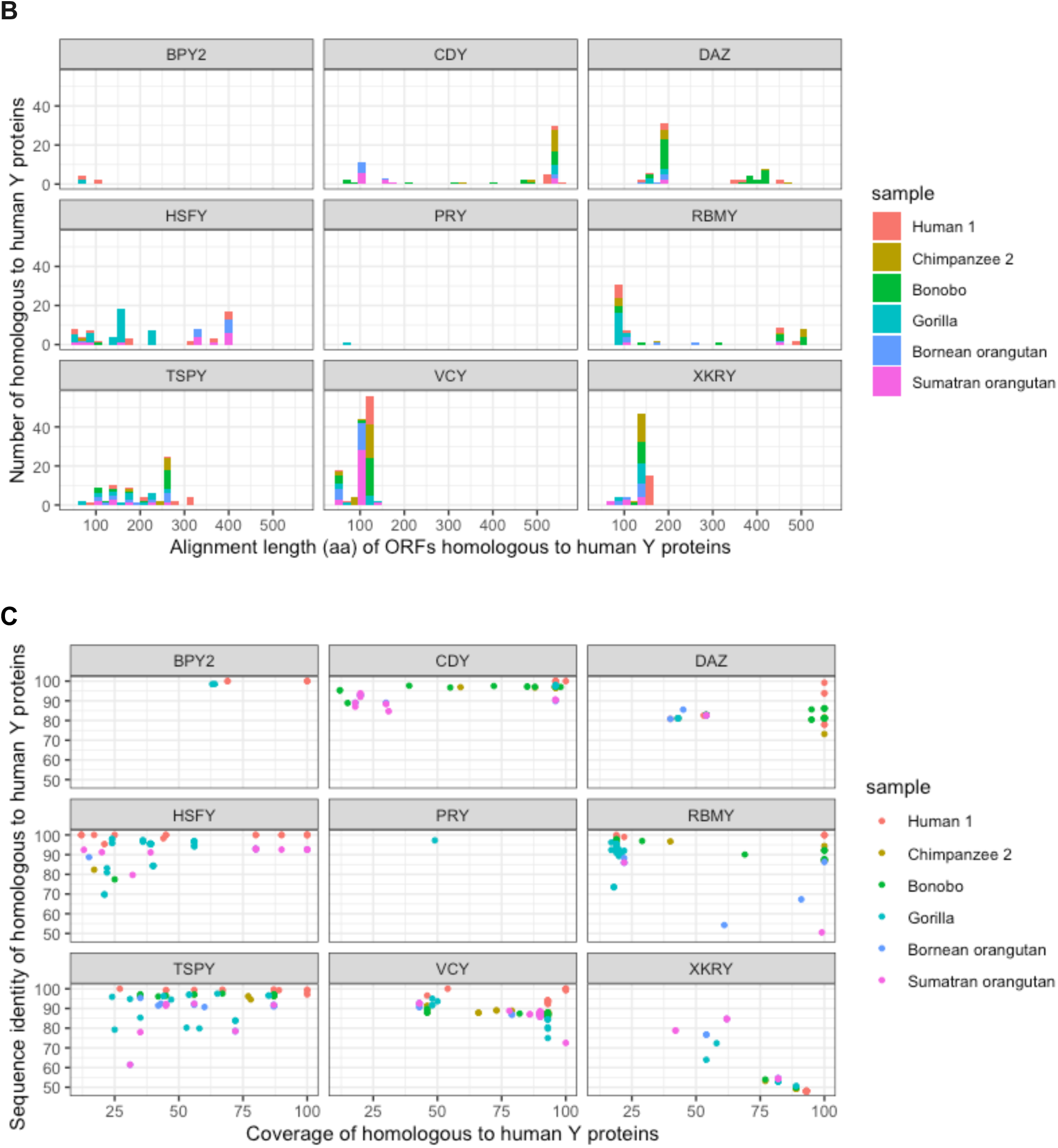
**A.** Analysis of high-confidence cORFs homologous to human Y-chromosome ampliconic transcripts per gene family per species. **A.** Numbers of cORFs. **B.** Alignment length (in amino acids) to human Y proteins. **C.** Coverage and sequence identity in alignments to homologous human Y proteins.

Predicted from cORFs CDY, DAZ, RBMY and TSPY proteins in chimpanzee and bonobo had high coverage and sequence identity when aligned to human Y proteins (Fig. 3C). Across great apes, the highest protein alignment length was observed for the CDY cORF homologs, followed by RBMY and DAZ cORF homologs (Fig. 3C). HSFY cORF homologs in Sumatran orangutan had high sequence identity to, and covered most of, the corresponding human Y proteins (Fig. 3C). Short HSFY cORF homologs of 50-250 aa were found in gorilla samples and longer ones (300-400 aa) were observed in both species of orangutan. Most VCY and XKRY cORF homologs from non-human great apes covered ~100-150 aa of the corresponding human Y proteins (Fig. 3C).

### Transcripts with identical protein-coding sequences

To gain insights into evolutionary conservation or divergence of YAG transcripts across great apes, we identified replicate-supported transcripts with identical cORFs (but potentially differing at 5’ and/or 3’ UTRs) within (between the two samples for human or chimpanzee) or between great ape species analyzed (Table S13, Dataset 14). We found that all gene families had shared cORFs between the two closely related orangutan species (Table S13). In contrast, some gene families (*BPY2* and *VCY*) consistently did not have shared cORFs between closely related Pan species, chimpanzee and bonobo (Table S13).In the whole dataset, most shared cORFs were present in *DAZ*, *PRY*, *RBMY,* and *TSPY* gene families (Table S13). *DAZ* and *PRY* cORFs were shared among human, chimpanzee, bonobo, and gorilla, but not between any one of them and the two orangutans, suggesting that human-gorilla is the largest evolutionary distance at which they are conserved. Identical *TSPY* cORFs were present within the same species (human or chimpanzee) and between some species (human and chimpanzee, human and bonobo, chimpanzee and bonobo, and the two species of orangutan; Table S13).

### Species-specific protein-coding transcript sequences

Subsequently, we were able to identify species-specific transcripts with cORFs (Table S16) and species-specific protein-coding sequences (Table S14). Across most gene families, at least one protein-coding transcript per gene family was species-specific. The highest number of species-specific protein-coding transcripts was present in the *PRY* gene family: it ranged from 16 to 57 per species (Table S14). *BPY2* had the lowest number of species-specific protein-coding transcripts, including three, three, and two transcripts specific to gorilla, Bornean orangutan, and Sumatran orangutan, respectively.

### Pseudogenes

Additionally, we identified many YAG transcripts without any cORFs, i.e. pseudogenes (Table S15). Most of them were present in the *PRY* gene family (min=9, max=85, and med=37 across species). The lowest number of pseudogenes was observed in the *XKRY*, *BPY2,* and *CDY* gene families (med=0 and maximum of 1, 2, and 2, respectively, across species).

### Alternative splicing patterns

To study splicing patterns of YAG transcripts across great apes, we mapped cORF-containing replicate-supported transcripts to one ampliconic gene copy (picked up at random) per gene family on the human Y chromosome (Figs. 4–5, S4–S11 showing only transcripts with cORFs, Datasets 15 and 16).

**Figure 4.**
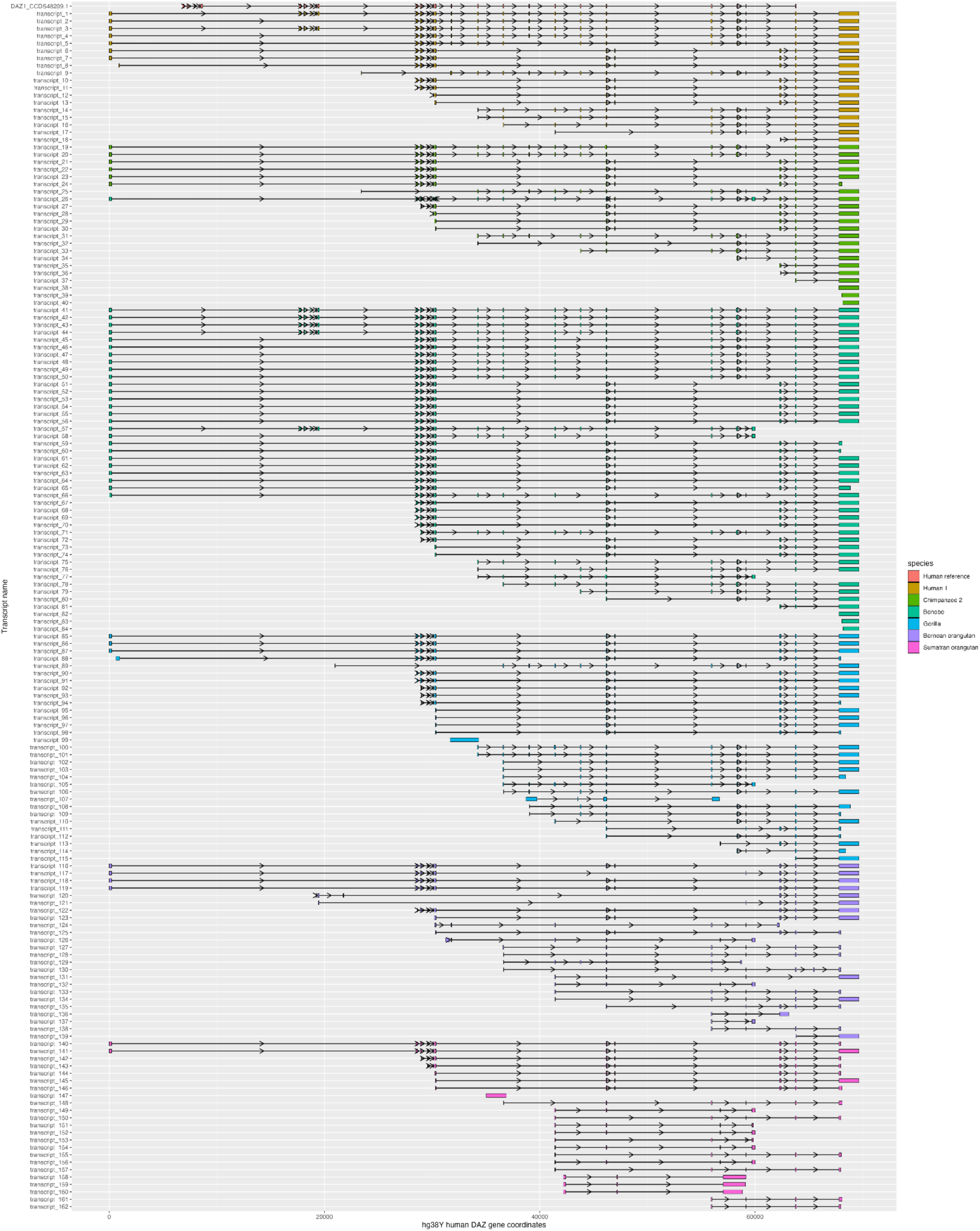
Conserved splicing patterns of *DAZ* transcripts with complete ORFs captured with gene-specific hybridization probes and mapped to one gene copy ENSG00000188120 on the X-axis. The human *DAZ* consensus coding sequence DAZ_CCDS48209.1 was also mapped to show the protein-coding exons (top row). Transcripts are colored by species. Each colored block represents a location of an exon, and each arrow indicates the forward “>” or reverse “<” direction as mapping to the human genomic copy. The lines between exons represent introns.

**Figure 5.**
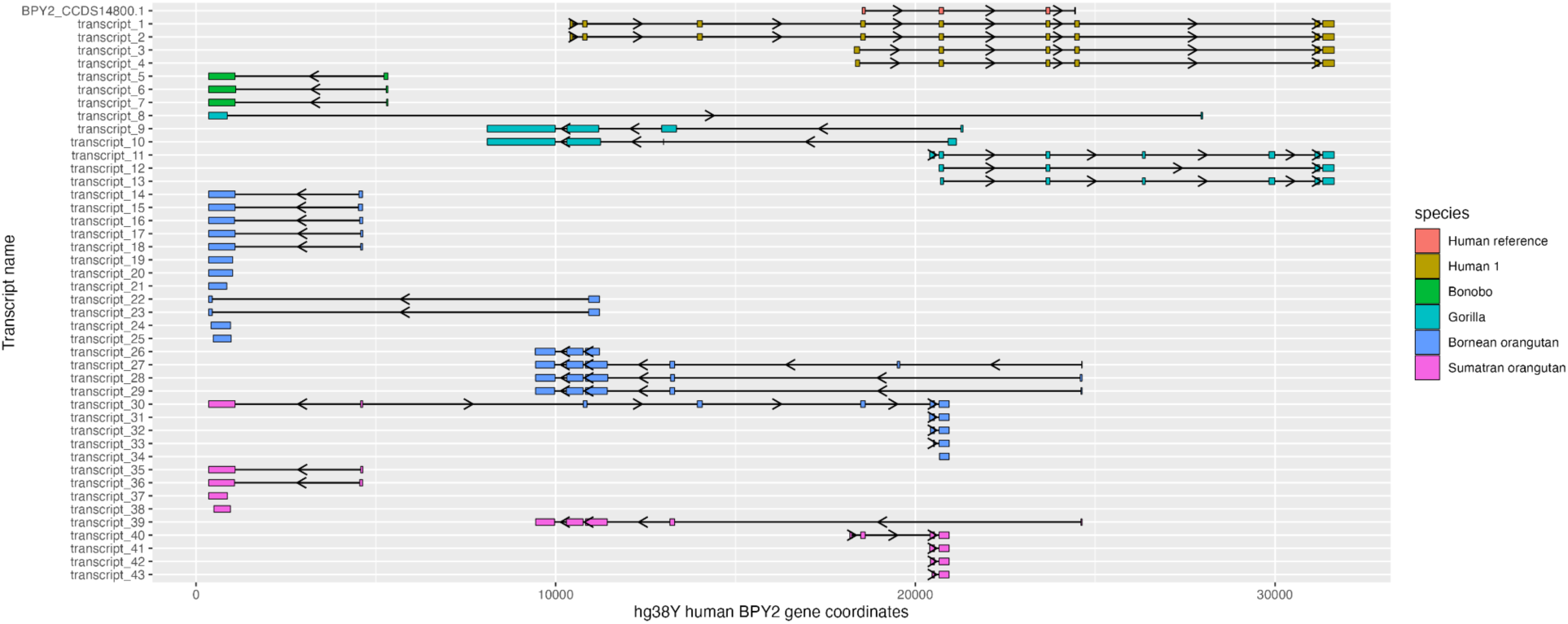
Divergent splicing patterns of *BPY2* transcripts with complete ORFs captured with gene-specific hybridization probes and mapped to one human gene copy ENSG00000183753. The human *BPY2* consensus coding sequence BPY2_CCDS14800.1 was also mapped to show the protein-coding exons (top row). Transcripts are colored by species. Each colored block represents a location of an exon, and each arrow indicates the forward “>” or reverse “<” direction as mapping to the human genomic copy. The lines between exons represent introns.

For some gene families, splicing patterns of non-human transcripts closely resembled that of human transcripts we recovered. Conserved splicing patterns are best exemplified by longest *DAZ* non-human transcripts, which, when mapped to the human genomic copy, had the exons’ locations similar to that of human transcripts (Fig. 4). Many non-human *CDY* transcripts recapitulated the splicing patterns of human transcripts (Fig. S4). Most non-human *HSFY* transcripts mapped to the same exon locations as did human transcripts (Fig. S5). Most non-human *RBMY* transcripts followed the locations of exons of human transcripts, with a few diverse transcripts found in gorilla and both species of orangutan (Fig. S7). Most non-human *TSPY* transcripts followed the splicing patterns of human transcripts (Fig. S8). *VCY* transcripts from bonobo and gorilla, which were captured with the human-specific probe, mapped to the exon locations of human transcripts (Fig. S9). *VCY* transcripts from chimpanzee and bonobo, captured with the Pan-specific probe, recapitulated the exon locations of chimpanzee transcripts (Fig. S10).

Other gene families had more divergent patterns of exon-intron structure among great apes. None of the exon locations for the *BPY2* transcripts from non-human samples recapitulated what was found in the human sample (Fig. 5). For example, many *BPY2* transcripts from bonobo and both species of orangutan mapped upstream from the human transcripts (Fig. 5). Additionally, all analyzed species but human and bonobo had *BPY2* transcripts originating from both strands of this gene (Fig. 5). All *PRY* transcripts from non-human samples mapped to only one exon of the human consensus coding sequence (Fig. S6). Most non-human *XKRY* transcripts mapped to the same two exons of human transcripts (Fig. S11), however all *XKRY* transcripts from both species of orangutan were missing the non-coding exon found in all the other species, and two *XKRY* transcripts from Bornean orangutan had longer protein-coding exonic sequences (Fig. S11).

### Selection tests

To test for selection, we limited our analysis to gene families that were present in five or more ape species as well as in the outgroup species (macaque): *CDY*, *DAZ*, *HSFY*, *RBMY*, and *TSPY*. We did not detect any evidence of positive selection in these gene families, however we did identify lower ratios of nonsynonymous over synonymous substitution rates in *CDY* in macaque and *DAZ* in human (Table S9). Nevertheless, the values were not significantly lower than one, rejecting the hypothesis that these sequences evolved under purifying selection (Table S9).

## DISCUSSION

Alternative splicing is one of the major forces generating the diversity and driving evolution of phenotypes across the eukaryotic tree of life. In mammals, alternative splicing has been widely investigated across tissues (Merkin et al. 2012; Gueroussov et al. 2015; Zhang et al. 2017) and recently also across developmental stages (Mazin et al. 2021). With recent increases in accuracy and read length of sequencing technologies, transcriptomic studies are unraveling unprecedented levels of diversity and complexity (Zhang et al. 2017; Oikonomopoulos et al. 2020; De Paoli-Iseppi, Gleeson, and Clark 2021; Bayega et al. 2018; Kuang and Canzar 2018). Deciphering transcript sequences of highly similar copies from multicopy gene families, however, still poses a significant challenge. Here, we focused on solving this problem by studying highly similar transcripts from multicopy YAG families, and, for the first time uncovered multiple transcripts for each of nine multicopy YAG families for all but one (Tapanuli orangutan) extant non-human great ape species. Earlier studies attempted to decipher transcript sequences of great ape YAGs using short-read RNA-seq datasets or to predict their full-length sequences using whole-genome DNA sequencing data (Cortez et al. 2014; Vegesna et al. 2020). However, such efforts resulted in reconstruction of only one consensus transcript per gene family and, in many cases, even that representative sequence was incomplete. A more specialized approach using RT-PCR and long-read sequencing was recently applied to capture multiple YAG transcripts per gene family from two human testis samples (Tomaszkiewicz and Makova 2018). Clustering YAG transcripts from long reads was achieved using a novel computational method (IsoCon (Sahlin et al. 2018)) that is able to distinguish transcripts originating from separate copies of a YAG family, which can differ by just a few nucleotides or small insertions/deletions. Applying this method to two human testis samples, Sahlin and colleagues (Sahlin et al. 2018) uncovered many novel YAG transcripts, however, because the primers were designed in the first and last protein-coding exons, full-length polyadenylated transcripts could not be obtained. To overcome this limitation, here we used gene-family-specific hybridization capture probes and the IsoSeq protocol, to pull down UTR-containing YAG cDNAs for human and five other great ape species. Additionally, to gain higher confidence in the sequence accuracy of our transcripts, we generated and sequenced transcripts from two technical replicates per each sample and were able to identify transcripts supported by both replicates.

### Diversity of YAG transcripts across great apes

We recovered at least two transcript sequences per YAG family per species, except for *BPY2* in chimpanzee (Table S6). Overall, we observed high variability in the number of transcripts per gene family per species. The high number of transcripts (303-315, Table S6) observed in both species of orangutan echoes the high copy number of YAGs reported previously in these species (Table S12) (Vegesna et al. 2020). Similarly, the low number of transcripts (190, Table S6) in chimpanzee is consistent with the low YAG copy number previously reported for this species (Table S12) (Vegesna et al. 2020). According to previous studies (Tomaszkiewicz et al. 2016; Vegesna et al. 2020), gorilla displayed the highest (7x) increase in transcript abundance with respect to YAG copy number (Table S12) and also one of the highest numbers of species-specific YAG protein-coding sequences (Table S14) and pseudogenes (Table S15).

### Conserved and divergent patterns of alternative splicing of YAG isoforms across great apes

Most gene families displayed conserved alternative splicing patterns across great apes, especially the longest *DAZ*, the single-exon *CDY*, the two-exon *HSFY*, and the multi-exon *TSPY* and *RBMY*. Most of transcripts from these gene families were conserved in human, chimpanzee, and bonobo, but were more divergent in gorilla and both species of orangutan. These divergent transcripts might encode proteins with different functions in spermatogenesis, a possibility that will have to be explored in future studies. Most divergent exon-intron structures were observed in the *BPY2* and *PRY* gene families, whose peculiarities are discussed below. Both *BPY2*, whose origin was only recently discovered (Cao et al. 2015), and *PRY* were excluded from previous evolutionary studies (e.g. (Bhowmick, Satta, and Takahata 2007)) because of the lack of information on their detectable X-linked/autosomal copies and on their orthologs in other species.

### Independent evolutionary origins of the BPY2 gene family transcripts

Our study suggests that *BPY2* evolution is conserved in human and chimpanzee, but is extremely rapid in other great apes; a previous study showed that BPY2 was acquired on the Y chromosome in the common ancestor of great apes (Cao et al. 2015). A full-length *BPY2* transcript was present in one of the chimpanzee technical replicates, suggesting that low expression in that sample might have prevented us from capturing the transcript in the other replicate. Consistent with this, *BPY2* was reported to have the second-lowest expression levels among YAGs (after *PRY)* in a previous study focused on great ape ampliconic gene expression levels (Vegesna et al. 2020). Additionally, *BPY2* is the only YAG family expressed during postmeiotic sex chromosome repression (Lucotte et al. 2018), the stage that might not have been captured in this chimpanzee sample. Nevertheless, this full-length *BPY2* transcript with a cORF found in one chimpanzee technical replicate aligned with high identity and over a large proportion of length to the human reference *BPY2* transcript (Dataset 10). Similarly, *BPY2* transcripts in orangutans mapped to human transcripts. In contrast, all *BPY2* transcripts from bonobo mapped upstream of human transcripts, suggesting their distinct evolutionary origins. Approximately half of gorilla *BPY2* transcripts mapped upstream of human transcripts. Interestingly, all species but human and bonobo had *BPY2* transcripts originating from both strands of the gene.

Our analysis of protein-coding regions suggested that BPY2 cORFs in several species might produce proteins nonhomologous to human proteins. Protein alignments between the predicted cORFs in non-human great apes and human Y-chromosome proteins produced no significant matches for *BPY2* sequences in bonobo and both species of orangutan (Fig. 3, Table S11). Gorilla BPY2 sequences had low identity and coverage when mapped to human Y BPY2 proteins. However, the recovered from one technical replicate chimpanzee BPY2 cORF had high (98%) sequence similarity over the full length (106 aa) human BPY2 protein (Dataset 12).

### *PRY* gene family is undergoing pseudogenization across great apes

Several lines of evidence in our results suggest that the *PRY* gene family is undergoing pseudogenization in most great ape species. First, most transcripts without cORFs we recovered map to the *PRY* gene family (Table S15). Second, all species but gorilla lack cORF-containing homologs to human *PRY*. Third, several *PRY* cORFs predicted for great apes are not homologous to human (Table S11) but are lineage-specific (Table S14) and mapping upstream of the human *PRY* coding exons (Fig. S13). Consistent with these results, among YAGs, *PRY* has been previously reported to be expressed at the lowest level, if at all, in great apes (Vegesna et al. 2020).

### Future applications to functional genomics

Cellular functions of YAGs are poorly characterized and were investigated only at the gene but not at the transcript level. For example, previous studies demonstrated that deletions of Y chromosome regions containing certain YAGs, i.e. azoospermia (AZF) regions, can lead to spermatogenic impairment—e.g. spermatogenic arrest resulting in altered spermatozoa formation or a complete lack of sperm cells (azoospermia) (Kuroda-Kawaguchi et al. 2001; Repping et al. 2002). Some other studies attempted to correlate the copy number of specific YAG families with fertility levels, however this led to inconclusive results (Krausz, Giachini, and Forti 2010; Nickkholgh et al. 2010; Giachini et al. 2009). Here, we uncovered splicing variants of YAGs that in the future can be used to study differential sperm characteristics and male fertility phenotypes observed across great apes. Overall, our study provides an informative genomic resource of full-length YAG transcripts for future functional studies focusing on infertility phenotypes in humans and other great apes.

### Limitations of the study and future directions

Ampliconic genes have been previously reported to be expressed at low levels in testes (Fagerberg et al. 2014; Vegesna et al. 2019, 2020). Though our study captured the highest transcript diversity to date, some of the genes, such as *BPY2* and *PRY,* were nevertheless not fully recovered in all the species. Thus, it would be recommended in the future to target these specific genes at a higher sequencing depth.

We have used one genomic copy per gene family from the human Y chromosome for tracking the splicing patterns of transcripts from human and non-human great ape samples. This allowed us to recapitulate the general conserved or divergent splicing patterns per gene family across great apes. In the future, a more detailed analysis with precise mapping to each well-annotated copy per gene family on the great ape Y chromosomes will alleviate human-specific bias of our analysis (Chen et al. 2021) of YAGs across great apes.

There are several potential technical reasons why we were unable to find any evidence of positive selection for any of the gene families. First, we did not test for positive selection in *BPY2*, *PRY*, *VCY*, and *XKRY,* which were either represented only in three species or missing from the outgroup. Thus, we might lack power to find evidence of selection in these most rapidly evolving gene families. Second, even though the codeml software from the PAML package (Yang 2007) is usually used to estimate the nonsynonymous to synonymous substitution rate ratios of genes, it does not account for deletions and insertions in splicing variants. This methodological limitation calls for developing other selection test approaches that include gaps representing divergence states in the alignments.

Testis is composed of several cell types: mitotic spermatogonia, meiotic spermatocytes, and post-meiotic spermatids and somatic cell types, such as Sertoli and Leydig cells. Thus, capturing ampliconic gene transcripts from testis samples does not allow for cell-specific distinction among transcripts. Future studies focusing on identifying cell-type specific YAG transcripts using single-cell RNA-seq approaches should overcome this limitation. Several recent studies investigated the expression levels of X-versus Y-linked genes including ampliconic genes in separate cell types from human testis, but none of them focused on analyzing the expression at the isoform level (Sin et al. 2012; Lucotte et al. 2018). One of the most recent studies has focused on single-nucleus testis transcriptome data from 11 species including four great apes (Murat et al. 2022). However, except for one *RBMY2* transcript from gorilla, no ampliconic transcripts were reported as a cell-type marker for any of the analyzed species. Additionally, it would be of interest to confirm all the transcripts we discovered in our study at the protein level, as was done in (Ferrández-Peral et al. 2022).

### Data access

PacBio IsoSeq data for each of the samples were submitted to the NCBI BioProject (http://www.ncbi.nlm.nih.gov/bioproject) under accession number PRJNA911852. Code is available at GitHub: https://github.com/makovalab-psu/YAG_analysis.

## ACKNOWLEDGEMENTS

We would like to thank Craig Praul and Dan Hannon of the Penn State Genomics Core Facility for PacBio sequencing. We are grateful to Andrew Cartoceti from the Smithsonian Institution, Oliver Ryder from the San Diego Zoo Institute for Conservation Research, and Sarah Dysart from MD Anderson Cancer Center for providing samples for this study. We thank Barbara Arbeithuber for early discussions on hybridization capture experiments, Bob Harris for providing us with the script for counting identical fasta sequences, and Barb McGrath for her critical reading of the manuscript. This study was funded by NIH Grant R01GM130691 (to K.D.M.), NIH Grant R01GM146462 (to P.M.) and NSF Grant DBI-2138585 (to P.M.).

## Supplementary Figures

**Figure S1.**
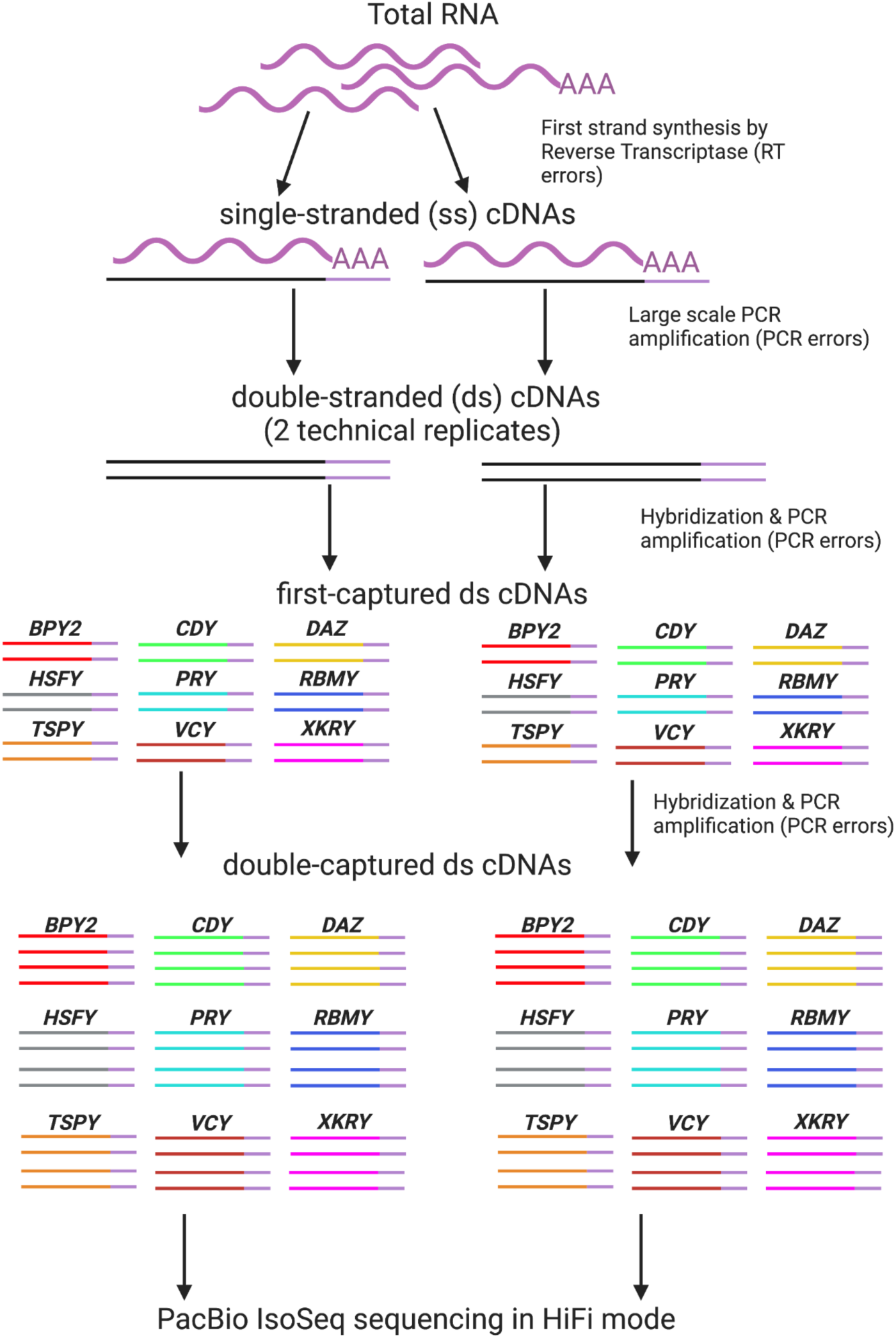
Experimental design. For each RNA sample, we generated two technical replicates started with cDNA synthesis and proceeded with the subsequent procedures in parallel. Two rounds of hybridization capture using the designed in-house capture probes (Table S2) resulted in gene-family-specific cDNAs, which were sequenced using PacBio HiFi sequencing. Generation of two technical replicates permits minimizes sequencing errors that could have been introduced in each of the technical replicates at random.

**Figure S2.**
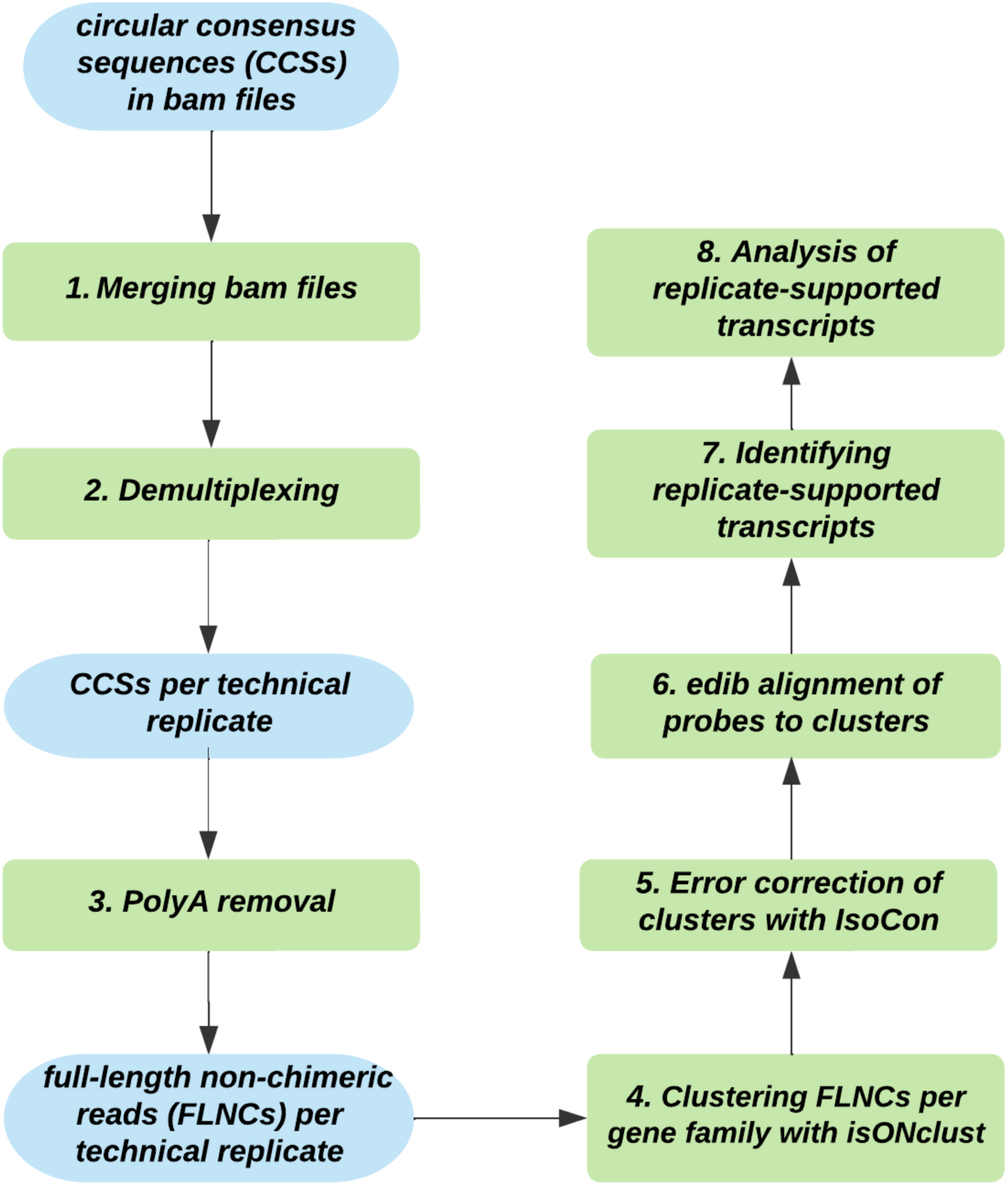
Workflow for the identification of replicate-supported shared transcripts.

**Figure S3.**
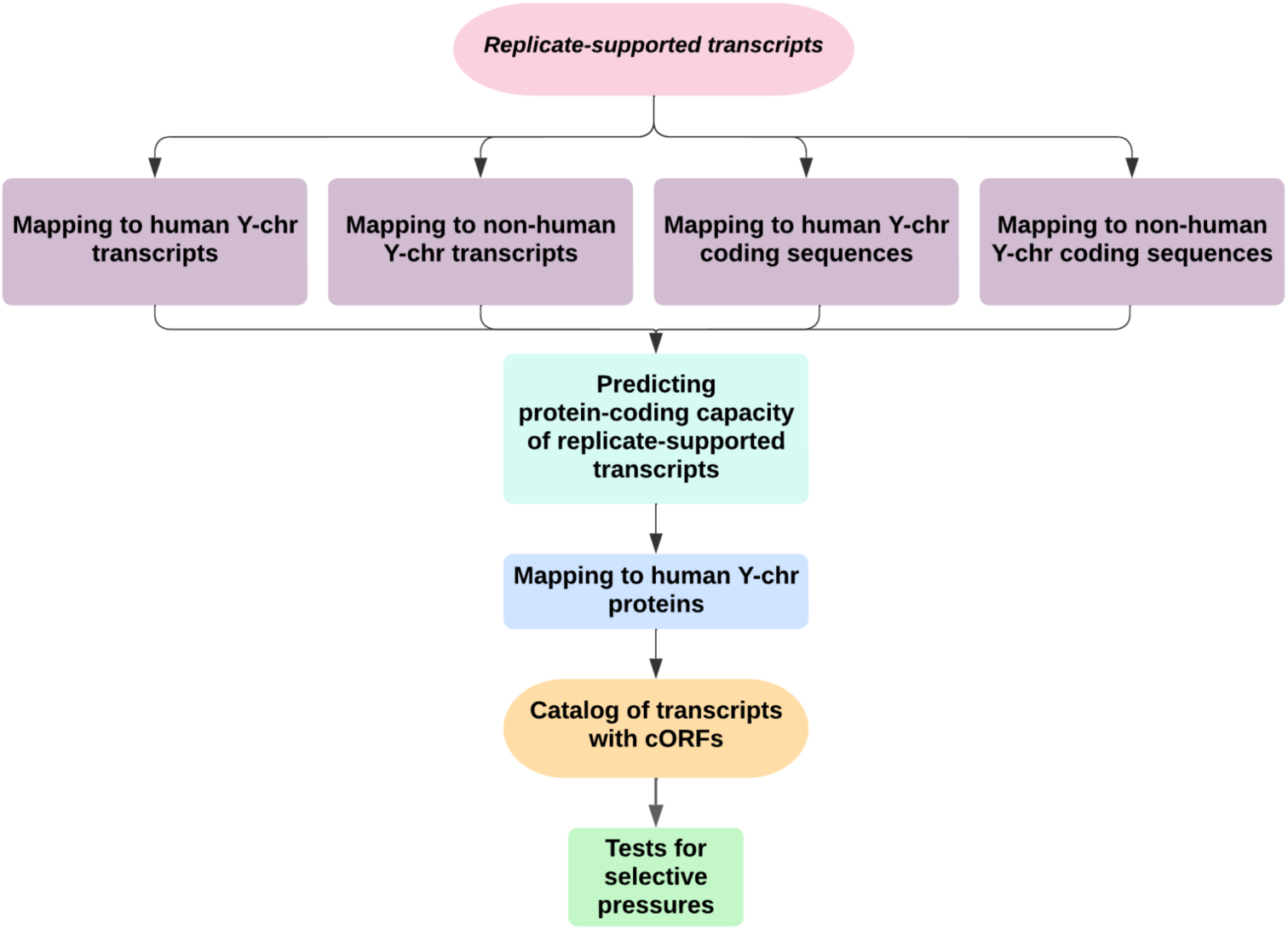
Flowchart for the biological analysis of replicate-supported transcripts.

**Figure S4.**
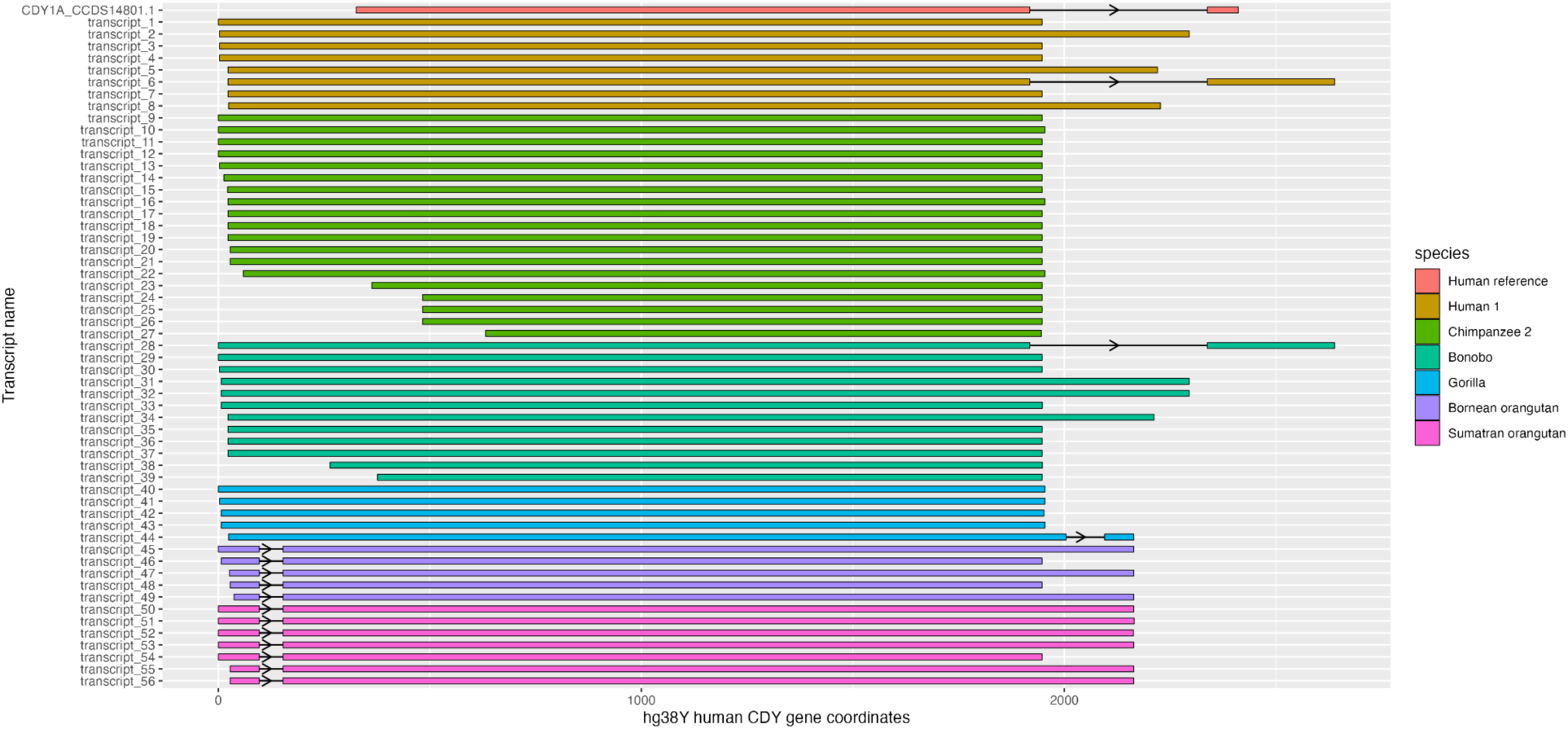
Splicing patterns of *CDY* transcripts with complete ORFs mapped to the location of one human gene copy ENSG00000172288. The human consensus coding sequence CDY1A_CCDS14801.1 was also mapped to show the protein-coding exons (top row). Transcripts are colored by species. Each colored block represents a location of an exon, and each arrow indicates the forward “>” or reverse “<” direction as mapped to the human genomic copy. The lines between exons represent introns.

**Figure S5.**
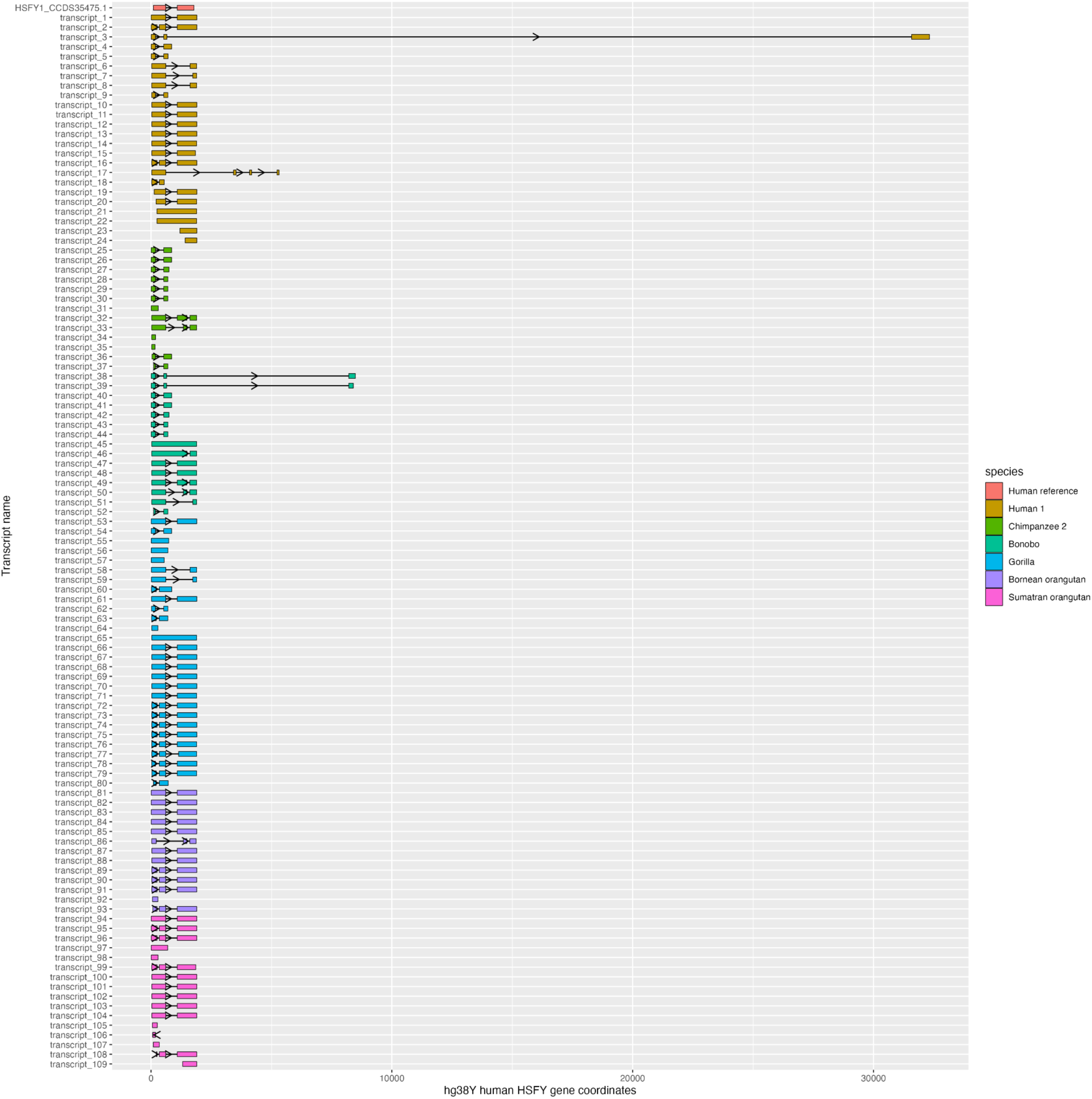
Splicing patterns of *HSFY* transcripts with complete ORFs mapped to one gene copy ENSG00000172468 on the X-axis. The human consensus coding sequence HSFY1_CCDS35475.1 was also mapped to show the protein-coding exons (top row). Transcripts are colored by species. Each colored block represents a location of an exon, and each arrow indicates the forward “>” or reverse “<” direction as mapping to the human genomic copy. The lines between exons represent introns.

**Figure S6.**
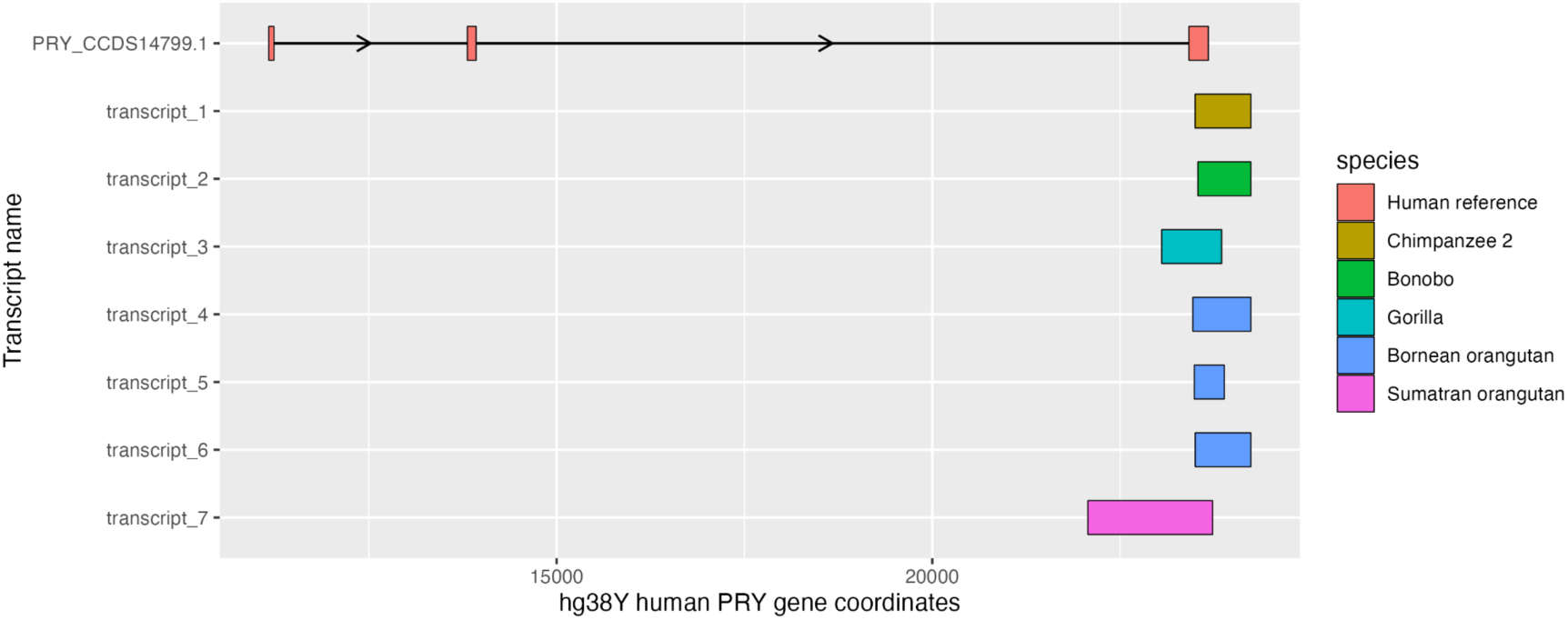
Splicing patterns of *PRY* transcripts with complete ORFs mapped to one gene copy ENSG00000169789 on the X-axis. The human *PRY* consensus coding sequence PRY_CCDS14799.1 was also mapped to show the protein-coding exons (top row). Transcripts are colored by species. Each colored block represents a location of an exon, and each arrow indicates the forward “>” or reverse “<” direction as mapping to the human genomic copy. The lines between exons represent introns.

**Figure S7.**
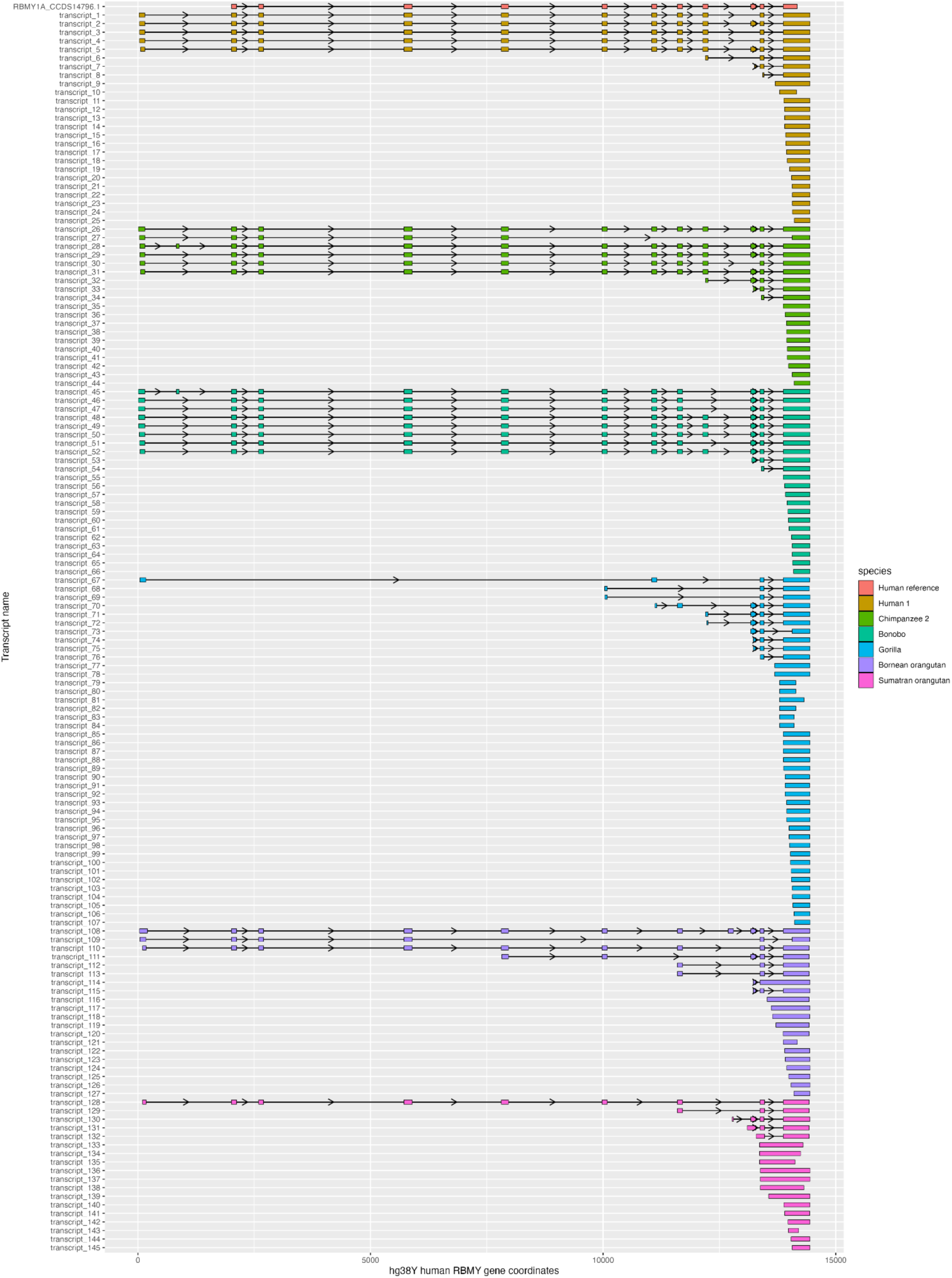
Splicing patterns of *RBMY* transcripts with complete ORFs mapped to one gene copy ENSG00000234414 on the X-axis. The human consensus coding sequence RBMY1A_CCDS14796.1 was also mapped to show the protein-coding exons (top row). Transcripts are colored by species. Each colored block represents a location of an exon, and each arrow indicates the forward “>” or reverse “<” direction as mapping to the human genomic copy. The lines between exons represent introns.

**Figure S8.**
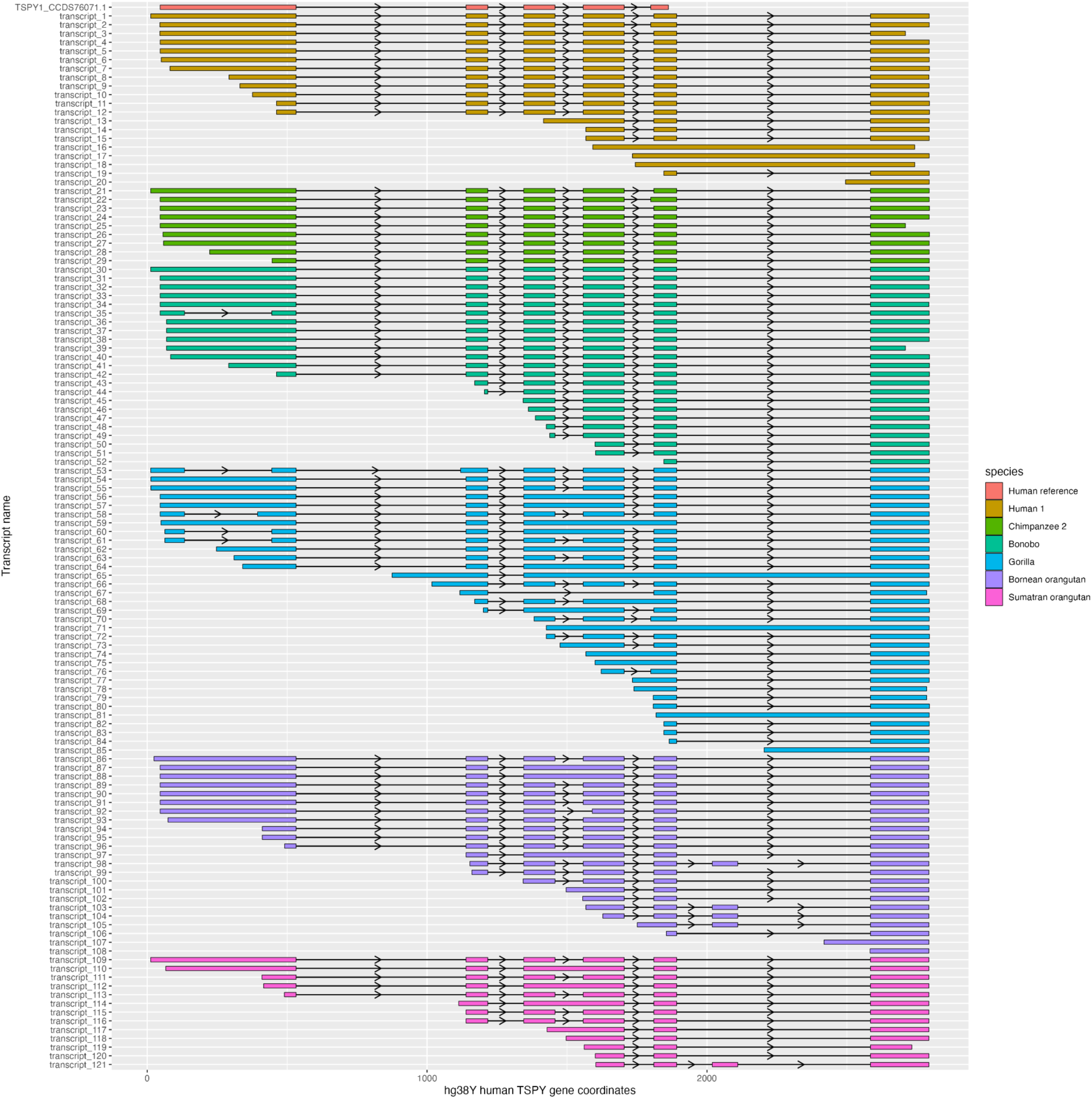
Splicing patterns of *TSPY* transcripts with complete ORFs mapped to one gene copy ENSG00000258992 on the X-axis. The human consensus coding sequence TSPY1_CCDS76071.1 was also mapped to show the protein-coding exons (top row). Transcripts are colored by species. Each colored block represents a location of an exon, and each arrow indicates the forward “>” or reverse “<” direction as mapping to the human genomic copy. The lines between exons represent introns.

**Figure S9.**
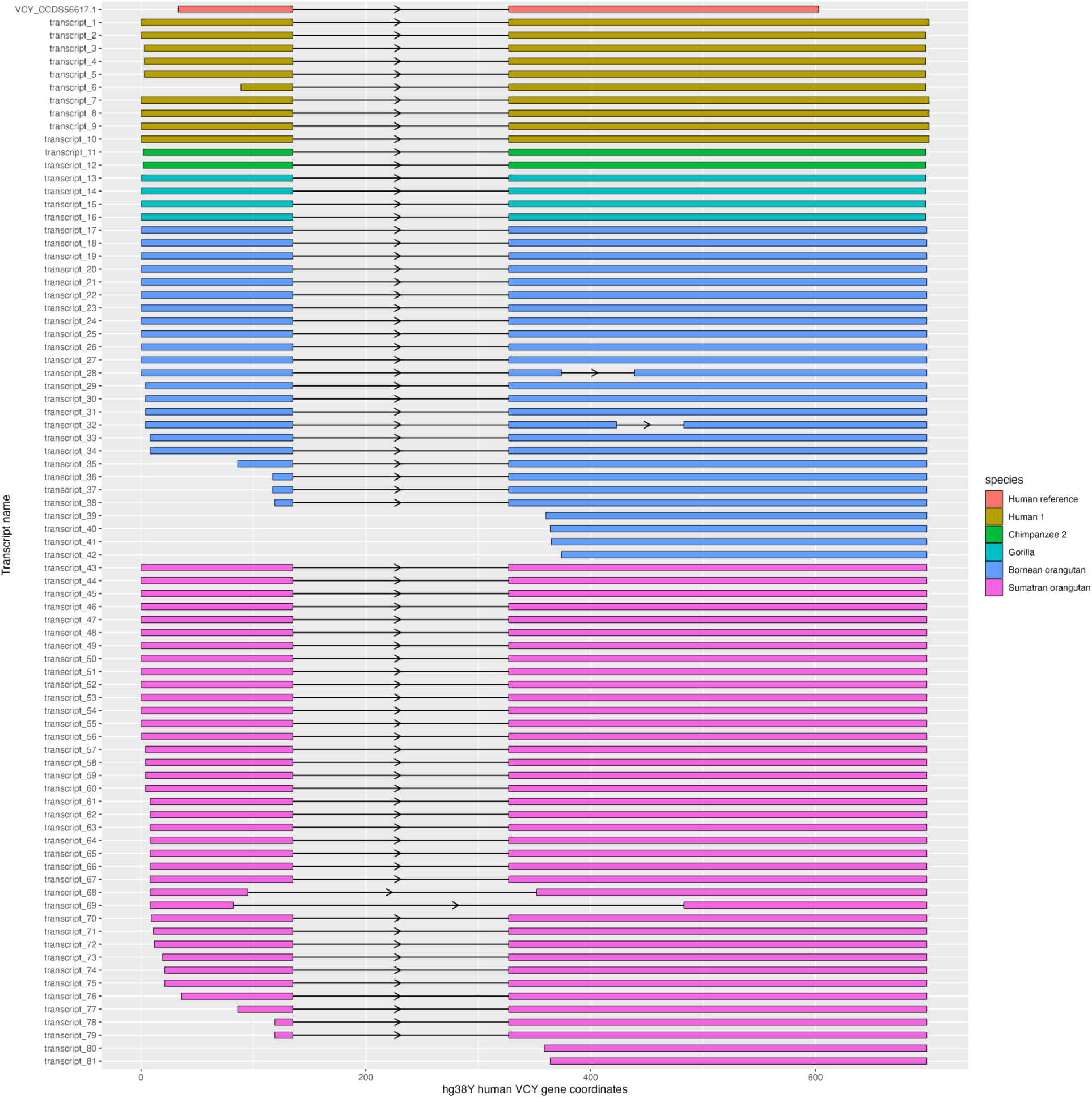
Splicing patterns of *VCY* transcripts with complete ORFs captured with *VCY*-human-specific hybridization probes and mapped to one human gene copy ENSG00000129864 on the X-axis. The human consensus coding sequence VCY_CCDS56617.1 was also mapped to show the protein-coding exons (top row). Transcripts are colored by species. Each colored block represents a location of an exon, and each arrow indicates the forward “>” or reverse “<” direction as mapping to the human genomic copy. The lines between exons represent introns.

**Figure S10.**
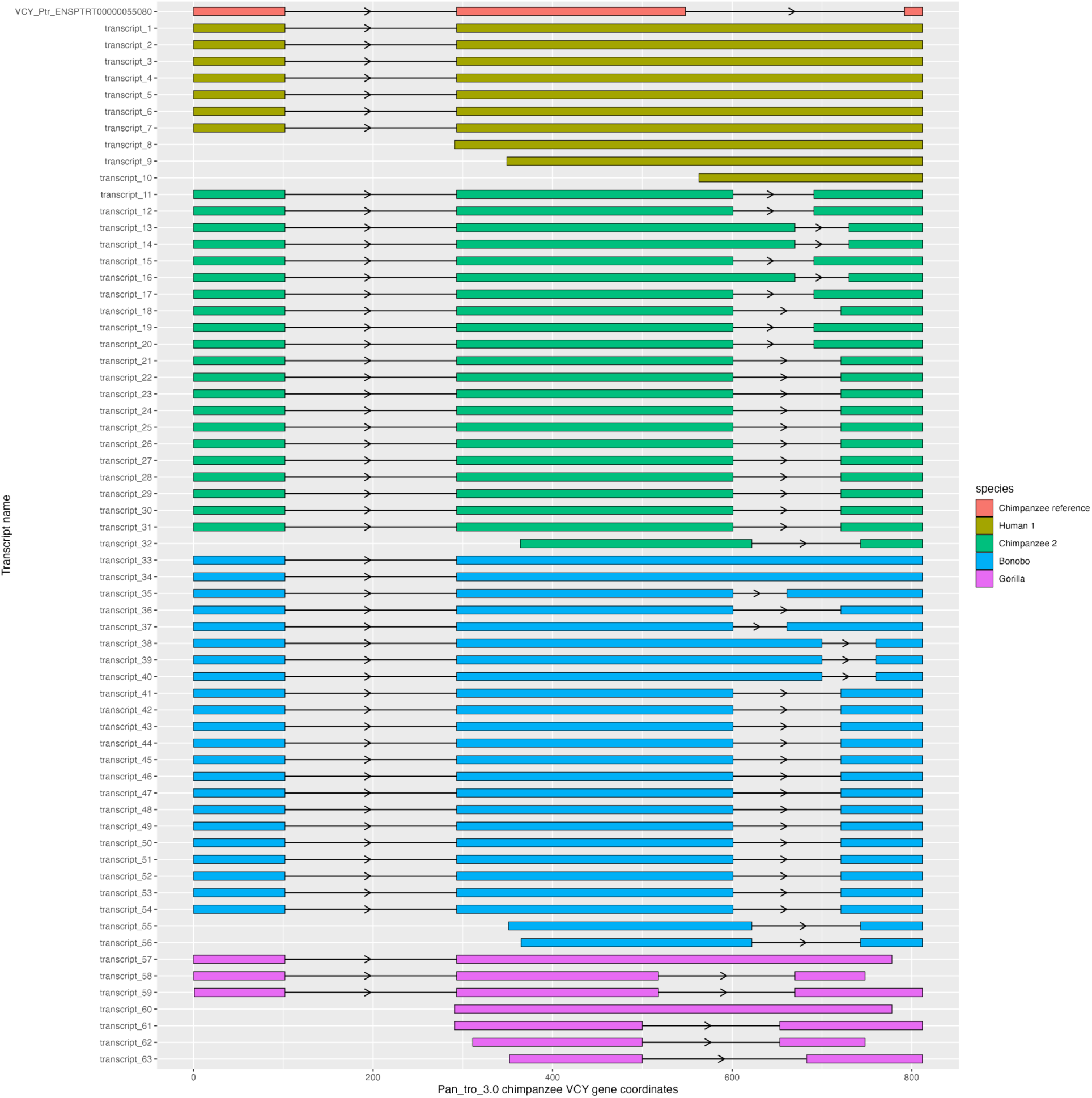
Splicing patterns of *VCY* transcripts with complete ORFs captured with *VCY*-Pan-specific hybridization probes and mapped to one chimpanzee gene copy ENSPTRG00000049568 on the X-axis. The chimpanzee *VCY* transcript sequence VCY_Ptr_ENSPTRT00000055080 was also mapped to show the protein-coding exons (top row). Transcripts are colored by species. Each colored block represents a location of an exon, and each arrow indicates the forward “>” or reverse “<” direction as mapping to the human genomic copy. The lines between exons represent introns. Some variation in alignments might be due to repeats present in the chimpanzee *VCY* gene copy ENSPTRG00000049568.

**Figure S11.**
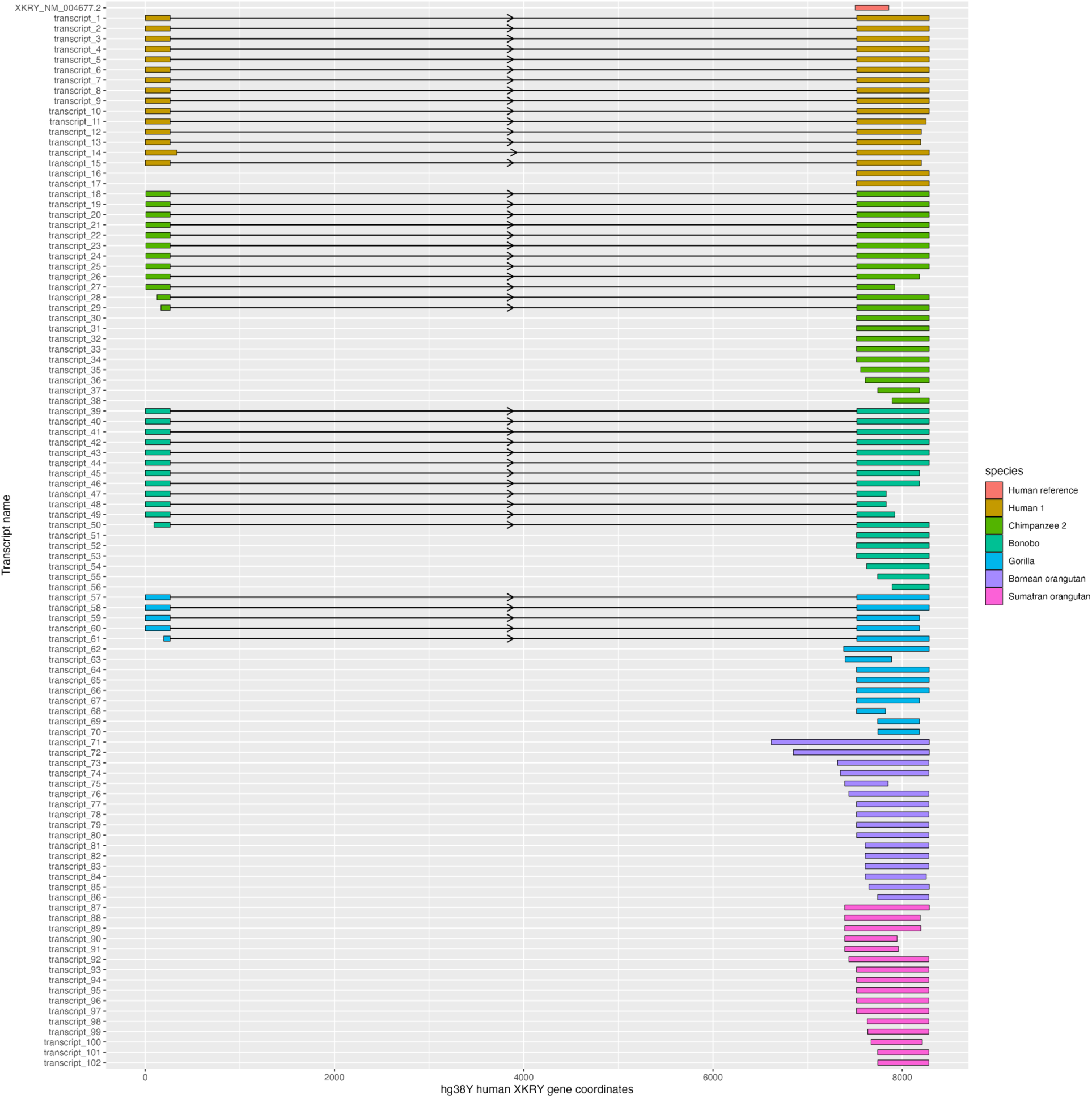
Splicing patterns of *XKRY* transcripts with complete ORFs captured with gene-specific hybridization probes and mapped to one gene copy ENSG00000250868 on the X-axis. The human *XKRY* transcript XKRY_NM_004677.2 was also mapped to show the protein-coding exons (top row). Transcripts are colored by species. Each colored block represents a location of an exon, and each arrow indicates the forward “>” or reverse “<” direction as mapping to the human genomic copy. The lines between exons represent introns.

## Notes

### Competing Interest Statement

The authors have declared no competing interest.

### Summary of Updates

Authorship and affiliations updated

## REFERENCES

Bayega, Anthony, Yu Chang Wang, Spyros Oikonomopoulos, Haig Djambazian, Somayyeh Fahiminiya, and Jiannis Ragoussis. 2018. “Transcript Profiling Using Long-Read Sequencing Technologies.” Methods in Molecular Biology 1783: 121–47.

Bellott, Daniel W., Jennifer F. Hughes, Helen Skaletsky, Laura G. Brown, Tatyana Pyntikova, Ting-Jan Cho, Natalia Koutseva, et al. 2014. “Mammalian Y Chromosomes Retain Widely Expressed Dosage-Sensitive Regulators.” Nature 508 (7497): 494–99.

Bhowmick, Bejon Kumar, Yoko Satta, and Naoyuki Takahata. 2007. “The Origin and Evolution of Human Ampliconic Gene Families and Ampliconic Structure.” Genome Research 17 (4): 441–50.

Camacho, Christiam, George Coulouris, Vahram Avagyan, Ning Ma, Jason Papadopoulos, Kevin Bealer, and Thomas L. Madden. 2009. “BLAST : Architecture and Applications.” BMC Bioinformatics. https://doi.org/10.1186/1471-2105-10-421.

Cao, Peng-Rong, Lei Wang, Yu-Chao Jiang, Yin-Sha Yi, Fang Qu, Tao-Cheng Liu, and Yuan Lv. 2015. “De Novo Origin of VCY2 from Autosome to Y-Transposed Amplicon.” PloS One 10 (3): e0119651.

Cechova, Monika, Rahulsimham Vegesna, Marta Tomaszkiewicz, Robert S. Harris, Di Chen, Samarth Rangavittal, Paul Medvedev, and Kateryna D. Makova. 2020. “Dynamic Evolution of Great Ape Y Chromosomes.” Proceedings of the National Academy of Sciences of the United States of America 117 (42): 26273–80.

Chen, Nae-Chyun, Brad Solomon, Taher Mun, Sheila Iyer, and Ben Langmead. 2021. “Reference Flow: Reducing Reference Bias Using Multiple Population Genomes.” Genome Biology 22 (1): 8.

Cortez, Diego, Ray Marin, Deborah Toledo-Flores, Laure Froidevaux, Angélica Liechti, Paul D. Waters, Frank Grützner, and Henrik Kaessmann. 2014. “Origins and Functional Evolution of Y Chromosomes across Mammals.” Nature 508 (7497): 488–93.

De Paoli-Iseppi, Ricardo, Josie Gleeson, and Michael B. Clark. 2021. “Isoform Age - Splice Isoform Profiling Using Long-Read Technologies.” Frontiers in Molecular Biosciences 8 (August): 711733.

Fagerberg, Linn, Björn M. Hallström, Per Oksvold, Caroline Kampf, Dijana Djureinovic, Jacob Odeberg, Masato Habuka, et al. 2014. “Analysis of the Human Tissue-Specific Expression by Genome-Wide Integration of Transcriptomics and Antibody-Based Proteomics.” Molecular & Cellular Proteomics: MCP 13 (2): 397–406.

Ferrández-Peral, Luis, Xiaoyu Zhan, Marina Alvarez-Estape, Cristina Chiva, Paula Esteller-Cucala, Raquel García-Pérez, Eva Julià, et al. 2022. “Transcriptome Innovations in Primates Revealed by Single-Molecule Long-Read Sequencing.” Genome Research 32 (8): 1448–62.

Frith, Martin C., Alistair R. Forrest, Ehsan Nourbakhsh, Ken C. Pang, Chikatoshi Kai, Jun Kawai, Piero Carninci, Yoshihide Hayashizaki, Timothy L. Bailey, and Sean M. Grimmond. 2006. “The Abundance of Short Proteins in the Mammalian Proteome.” PLoS Genetics. https://doi.org/10.1371/journal.pgen.0020052.

Giachini, Claudia, Francesca Nuti, Daniel J. Turner, Ilaria Laface, Yali Xue, Fabrice Daguin, Gianni Forti, Chris Tyler-Smith, and Csilla Krausz. 2009. “TSPY1 Copy Number Variation Influences Spermatogenesis and Shows Differences among Y Lineages.” The Journal of Clinical Endocrinology & Metabolism. https://doi.org/10.1210/jc.2009-1029.

Glazko, G. V. 2003. “Estimation of Divergence Times for Major Lineages of Primate Species.” Molecular Biology and Evolution. https://doi.org/10.1093/molbev/msg050.

Gueroussov, Serge, Thomas Gonatopoulos-Pournatzis, Manuel Irimia, Bushra Raj, Zhen-Yuan Lin, Anne-Claude Gingras, and Benjamin J. Blencowe. 2015. “An Alternative Splicing Event Amplifies Evolutionary Differences between Vertebrates.” Science 349 (6250): 868–73.

Hughes, Jennifer F., Helen Skaletsky, Tatyana Pyntikova, Tina A. Graves, Saskia K. M. van Daalen, Patrick J. Minx, Robert S. Fulton, et al. 2010. “Chimpanzee and Human Y Chromosomes Are Remarkably Divergent in Structure and Gene Content.” Nature 463 (7280): 536–39.

Krausz, Csilla, Claudia Giachini, and Gianni Forti. 2010. “TSPY and Male Fertility.” Genes 1 (2): 308–16.

Kuang, Zheng, and Stefan Canzar. 2018. “Tracking Alternatively Spliced Isoforms from Long Reads by SpliceHunter.” Methods in Molecular Biology 1751: 73–88.

Kumar, Sudhir, Glen Stecher, Michael Li, Christina Knyaz, and Koichiro Tamura. 2018. “MEGA X: Molecular Evolutionary Genetics Analysis across Computing Platforms.” Molecular Biology and Evolution 35 (6): 1547–49.

Kuroda-Kawaguchi, Tomoko, Helen Skaletsky, Laura G. Brown, Patrick J. Minx, Holland S. Cordum, Robert H. Waterston, Richard K. Wilson, et al. 2001. “The AZFc Region of the Y Chromosome Features Massive Palindromes and Uniform Recurrent Deletions in Infertile Men.” Nature Genetics. https://doi.org/10.1038/ng757.

Lahn, B. T., and D. C. Page. 1997. “Functional Coherence of the Human Y Chromosome.” Science 278 (5338): 675–80.

Lahn, B. T., and D. C. Page. 2000. “A Human Sex-Chromosomal Gene Family Expressed in Male Germ Cells and Encoding Variably Charged Proteins.” Human Molecular Genetics 9 (2): 311–19.

Larkin, M. A., G. Blackshields, N. P. Brown, R. Chenna, P. A. McGettigan, H. McWilliam, F. Valentin, et al. 2007. “Clustal W and Clustal X Version 2.0.” Bioinformatics. https://doi.org/10.1093/bioinformatics/btm404.

Li, Heng. 2013. “Aligning Sequence Reads, Clone Sequences and Assembly Contigs with BWA-MEM.” *arXiv*:1303.3997 [q-bio.GN]. https://doi.org/https://arxiv.org/abs/1303.3997.

Locke, Devin P., Ladeana W. Hillier, Wesley C. Warren, Kim C. Worley, Lynne V. Nazareth, Donna M. Muzny, Shiaw-Pyng Yang, et al. 2011. “Comparative and Demographic Analysis of Orang-Utan Genomes.” Nature 469 (7331): 529–33.

Lucotte, Elise A., Laurits Skov, Jacob Malte Jensen, Moisès Coll Macià, Kasper Munch, and Mikkel H. Schierup. 2018. “Dynamic Copy Number Evolution of X- and Y-Linked Ampliconic Genes in Human Populations.” Genetics 209 (3): 907–20.

Mazin, Pavel V., Philipp Khaitovich, Margarida Cardoso-Moreira, and Henrik Kaessmann. 2021. “Alternative Splicing during Mammalian Organ Development.” Nature Genetics 53 (6): 925–34.

Merkin, Jason, Caitlin Russell, Ping Chen, and Christopher B. Burge. 2012. “Evolutionary Dynamics of Gene and Isoform Regulation in Mammalian Tissues.” Science 338 (6114): 1593–99.

Murat, Florent, Noe Mbengue, Sofia Boeg Winge, Timo Trefzer, Evgeny Leushkin, Mari Sepp, Margarida Cardoso-Moreira, et al. 2022. “The Molecular Evolution of Spermatogenesis across Mammals.” Nature, December. https://doi.org/10.1038/s41586-022-05547-7.

Navarro-Costa, Paulo. 2012. “Sex, Rebellion and Decadence: The Scandalous Evolutionary History of the Human Y Chromosome.” Biochimica et Biophysica Acta 1822 (12): 1851–63.

Nickkholgh, Bita, Michiel J. Noordam, Suzanne E. Hovingh, Ans M. M. van Pelt, Fulco van der Veen, and Sjoerd Repping. 2010. “Y Chromosome TSPY Copy Numbers and Semen Quality.” Fertility and Sterility 94 (5): 1744–47.

Oetjens, Matthew T., Feichen Shen, Sarah B. Emery, Zhengting Zou, and Jeffrey M. Kidd. 2016. “Y-Chromosome Structural Diversity in the Bonobo and Chimpanzee Lineages.” Genome Biology and Evolution 8 (7): 2231–40.

Oikonomopoulos, Spyros, Anthony Bayega, Somayyeh Fahiminiya, Haig Djambazian, Pierre Berube, and Jiannis Ragoussis. 2020. “Methodologies for Transcript Profiling Using Long-Read Technologies.” Frontiers in Genetics 11 (July): 606.

Pearson, William R. 2013. “An Introduction to Sequence Similarity (‘Homology’) Searching.” Current Protocols in Bioinformatics. https://doi.org/10.1002/0471250953.bi0301s42.

Repping, Sjoerd, Helen Skaletsky, Julian Lange, Sherman Silber, Fulco Van Der Veen, Robert D. Oates, David C. Page, and Steve Rozen. 2002. “Recombination between Palindromes P5 and P1 on the Human Y Chromosome Causes Massive Deletions and Spermatogenic Failure.” American Journal of Human Genetics 71 (4): 906–22.

Rice, P., I. Longden, and A. Bleasby. 2000. “EMBOSS: The European Molecular Biology Open Software Suite.” Trends in Genetics: TIG 16 (6): 276–77.

Sahlin, Kristoffer, and Veli Mäkinen. 2021. “Accurate Spliced Alignment of Long RNA Sequencing Reads.” Bioinformatics, July. https://doi.org/10.1093/bioinformatics/btab540.

Sahlin, Kristoffer, and Paul Medvedev. 2020. “De Novo Clustering of Long-Read Transcriptome Data Using a Greedy, Quality Value-Based Algorithm.” Journal of Computational Biology: A Journal of Computational Molecular Cell Biology 27 (4): 472–84.

Sahlin, Kristoffer, Marta Tomaszkiewicz, Kateryna D. Makova, and Paul Medvedev. 2018. “Deciphering Highly Similar Multigene Family Transcripts from Iso-Seq Data with IsoCon.” Nature Communications 9 (1): 4601.

Sin, Ho-Su, Yosuke Ichijima, Eitetsu Koh, Mikio Namiki, and Satoshi H. Namekawa. 2012. “Human Postmeiotic Sex Chromatin and Its Impact on Sex Chromosome Evolution.” Genome Research 22 (5): 827–36.

Skaletsky, Helen, Tomoko Kuroda-Kawaguchi, Patrick J. Minx, Holland S. Cordum, Ladeana Hillier, Laura G. Brown, Sjoerd Repping, et al. 2003. “The Male-Specific Region of the Human Y Chromosome Is a Mosaic of Discrete Sequence Classes.” Nature 423 (6942): 825–37.

Šošić, Martin, and Mile Šikić. 2017. “Edlib: A C/C ++ Library for Fast, Exact Sequence Alignment Using Edit Distance.” Bioinformatics 33 (9): 1394–95.

Stouffs, K. 2004. “Expression Pattern of the Y-Linked PRY Gene Suggests a Function in Apoptosis but Not in Spermatogenesis.” Molecular Human Reproduction. https://doi.org/10.1093/molehr/gah010.

Thorvaldsdóttir, Helga, James T. Robinson, and Jill P. Mesirov. 2013. “Integrative Genomics Viewer (IGV): High-Performance Genomics Data Visualization and Exploration.” Briefings in Bioinformatics 14 (2): 178–92.

Tomaszkiewicz, Marta, and Kateryna Makova. 2018. “Targeted Sequencing of Ampliconic Gene Transcripts from Total Human Male Testis RNA.” Protocol Exchange. https://doi.org/10.1038/protex.2018.109.

Tomaszkiewicz, Marta, Samarth Rangavittal, Monika Cechova, Rebeca Campos Sanchez, Howard W. Fescemyer, Robert Harris, Danling Ye, et al. 2016. “A Time- and Cost-Effective Strategy to Sequence Mammalian Y Chromosomes: An Application to the de Novo Assembly of Gorilla Y.” Genome Research 26 (4): 530–40.

Vegesna, Rahulsimham, Marta Tomaszkiewicz, Paul Medvedev, and Kateryna D. Makova. 2019. “Dosage Regulation, and Variation in Gene Expression and Copy Number of Human Y Chromosome Ampliconic Genes.” PLoS Genetics 15 (9): e1008369.

Vegesna, Rahulsimham, Marta Tomaszkiewicz, Oliver A. Ryder, Rebeca Campos-Sánchez, Paul Medvedev, Michael DeGiorgio, and Kateryna D. Makova. 2020. “Ampliconic Genes on the Great Ape Y Chromosomes: Rapid Evolution of Copy Number but Conservation of Expression Levels.” Genome Biology and Evolution 12 (6): 842–59.

Wong, Elaine Y. M., Jenny Y. M. Tse, Kwok-Ming Yao, Vincent C. H. Lui, Po-Chor Tam, and William S. B. Yeung. 2004. “Identification and Characterization of Human VCY2-Interacting Protein: VCY2IP-1, a Microtubule-Associated Protein-like Protein.” Biology of Reproduction 70 (3): 775–84.

Yang, Ziheng. 2007. “PAML 4: Phylogenetic Analysis by Maximum Likelihood.” Molecular Biology and Evolution 24 (8): 1586–91.

Ye, Danling, Arslan A. Zaidi, Marta Tomaszkiewicz, Kate Anthony, Corey Liebowitz, Michael DeGiorgio, Mark D. Shriver, and Kateryna D. Makova. 2018. “High Levels of Copy Number Variation of Ampliconic Genes across Major Human Y Haplogroups.” Genome Biology and Evolution. https://doi.org/10.1093/gbe/evy086.

Yen, Pauline H. 2004. “Putative Biological Functions of the DAZ Family.” International Journal of Andrology 27 (3): 125–29.

Zhang, Shi-Jian, Chenqu Wang, Shouyu Yan, Aisi Fu, Xuke Luan, Yumei Li, Qing Sunny Shen, et al. 2017. “Isoform Evolution in Primates through Independent Combination of Alternative RNA Processing Events.” Molecular Biology and Evolution 34 (10): 2453–68.

Zou, Sheng Wei, Jian Chao Zhang, Xiao Dong Zhang, Shi Ying Miao, Shu Dong Zong, Qi Sheng, and Lin Fang Wang. 2003. “Expression and Localization of VCX/Y Proteins and Their Possible Involvement in Regulation of Ribosome Assembly during Spermatogenesis.” Cell Research. https://doi.org/10.1038/sj.cr.7290161.

